# IKAP - Identifying K mAjor cell Population groups in single-cell RNA-seq analysis

**DOI:** 10.1101/596817

**Authors:** Yun-Ching Chen, Abhilash Suresh, Chingiz Underbayev, Clare Sun, Komudi Singh, Fayaz Seifuddin, Adrian Wiestner, Mehdi Pirooznia

## Abstract

In single-cell RNA-seq analysis, clustering cells into groups and differentiating cell groups by marker genes are two separate steps for investigating cell identity. However, results in clustering greatly affect the ability to differentiate between cell groups. We develop IKAP – an algorithm identifying major cell groups that improves differentiating by tuning parameters for clustering. Using multiple datasets, we demonstrate IKAP improves identification of major cell types and facilitates cell ontology curation.

## Main

Single-cell RNA-sequencing (scRNA-seq) enables inquiry of cell identity based on single cell transcriptomics. To facilitate cell type characterization and recognition, computational methods have been developed for (*i*) clustering cells with similar transcriptomic profiles into groups and (*ii*) identifying a set of marker genes to differentiate those cell groups (1). These two tasks have been treated independently although groups identified by clustering greatly determine the marker genes associated with each group. Because computing marker genes is often more resource intensive compared to clustering, we attempted to improve and accelerate biological interpretation by developing an algorithm to effectively identify the *k* major groups that produce distinguishing marker genes.

Despite the existence of well-performing scRNA clustering methods, identifying *k* groups remains challenging due to parameter specification (2). Most (if not all) clustering methods require a parameter suggesting *k* and a list of features such as genes or principal components (PCs) for computing cell-to-cell similarity. The proper *k* is generally unknown a priori. Choosing a small *k* may mix more than one cell type in a group whereas choosing a large *k* would result in many subgroups of unclear biological significance. Both can complicate cell type recognition by producing uninformative marker genes. In addition, the choice of feature list can affect grouping quality which, in turn, affects its distinguishing power. Therefore, *k* and feature list selection often become a bottleneck in the scRNA analysis pipeline.

To address this issue, we propose an unbiased approach – called IKAP (<u>I</u>dentifying <u>K</u> m<u>A</u>jor cell Population groups) – which identifies the *k* well-separated major groups likely to produce distinguishing marker genes in a scRNA dataset by systematically exploring the parameter space (Figure 1 and Online Methods). IKAP is implemented on top of Seurat (3) – one of the most widely used scRNA analysis packages – in which clustering requires two user-specified parameters: resolution *r* that determines *k* (the higher *r*, the larger *k*) and the number of top principal components (nPC) as the feature list. Briefly, for a given nPC, IKAP initializes a set of *k*_*max*_ groups by setting a high *r*. To simulate the coarse-to-fine grouping process, two nearest groups are merged iteratively, generating *k*_*max*_ sets of groups with *k* = 1 to *k*_*max*_. For each set, gap statistic is computed to measure the gap between the grouping with observed data and that with random data (4). The gap often monotonically increases (at variable amount) as *k* increases from 1 to *k*_*max*_ indicating that splitting out each group somewhat contributes to the grouping moving away from randomness (Supplementary Figure 1). We reason that those *k’s* that contribute more (*i.e*. yield large gap increase) might correspond to the set of *k* well-separated major groups. IKAP repeats this procedure for a range of nPCs. Finally, candidate sets grouped by clustering cells with different *k’s* and nPCs are picked from those with larger gap increase. Among all candidate sets, the one with the lowest classification error is marked as the best using decision trees built from marker genes. IKAP can be run by default without specifying any parameter as we did for all experiments in this study and can potentially be tailored for scRNA clustering methods other than Seurat.

**Figure 1.**
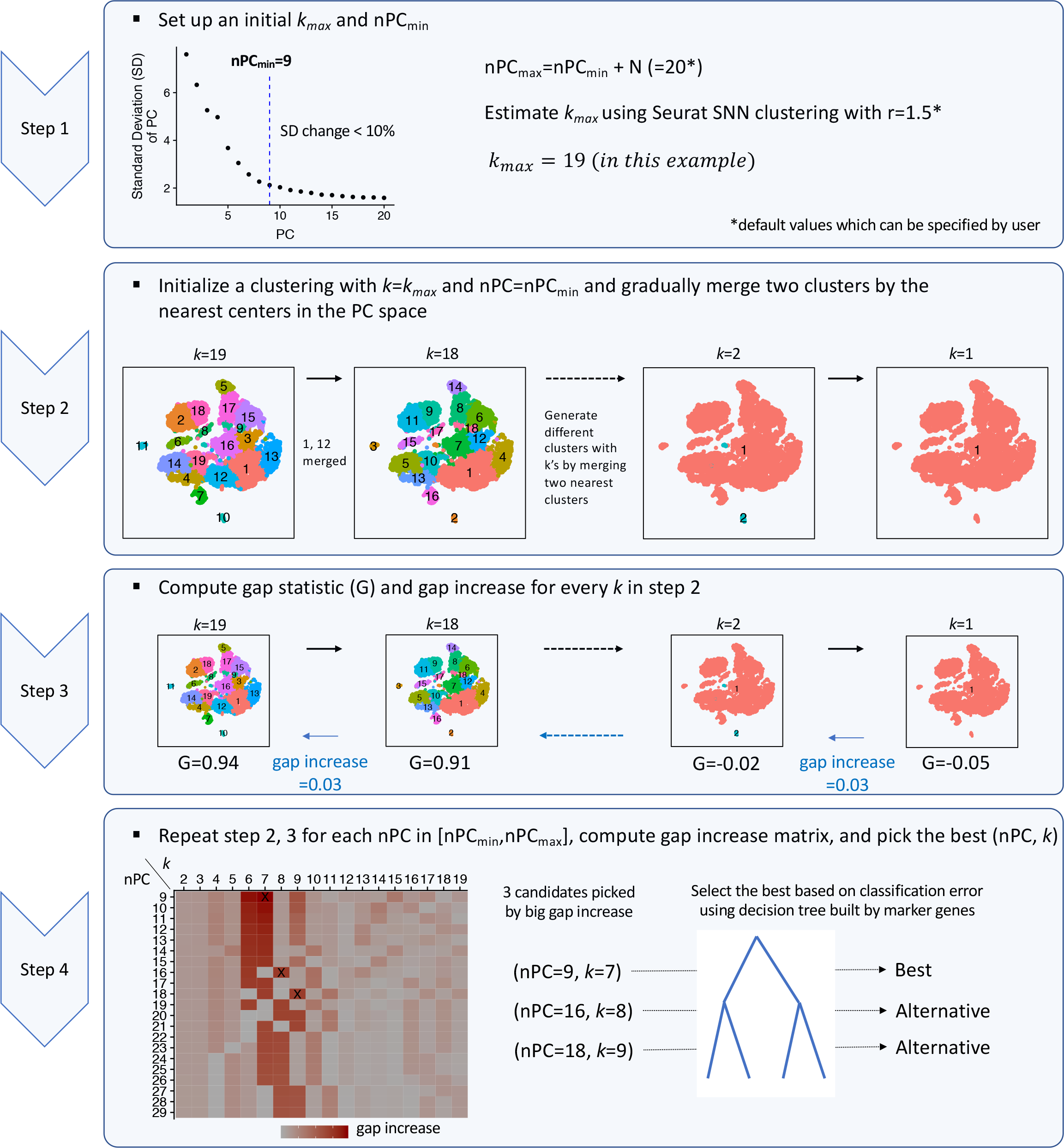
IKAP workflow. See Online Methods for details.

We tested IKAP on a peripheral blood mononuclear cell (PBMC) dataset of ~8K cells (denoted as PBMC_8K) from a healthy donor (5). The best set (with *k=7* and nPC=9; thus, abbreviated as PC9K7) and two alternative sets (PC16K8 and PC18K9 respectively) were reported. The major groups reported in PC9K7 were effectively aligned with known major cell lineages such as B cells, T cells, and NK cells as evidenced by expression of known marker genes (Figure 2A). These known marker genes of different cell lineages (such as *CD3E, TRAC*, and *IL32* for T cells) were also prioritized to the top of the marker gene list for every group (Figure 2B), facilitating cell type determination. To compare with the trial-and-error strategy, we varied nPC (=5, 10, 15, and 20) and *r* (=0.1, 0.2, 0.4, 0.6, and 1.0) to generate 20 trial sets of groups using Seurat clustering. Most trial sets did not divide cells into major cell types (Supplementary Figure 2) such that cell lineage marker genes were not ranked at the top or were unspecific to particular cell groups, complicating cell type recognition (Supplementary Figure 3). To quantitatively evaluate whether a set of cell groups can produce distinguishing marker genes, we designed three metrics: (*i*) the number of marker genes with high AUROC (Area Under the ROC curve), (*ii*) in-group versus out-of-group expression fold change among high AUROC marker genes, and (*iii*) classification error when classifying cells using decision trees built from multiple marker genes. Compared with the 20 trial sets, we found PC9K7 yielded more marker genes with high AUROC, higher expression fold change, and lower classification error (Figure 2C). Two alternative sets (PC16K8 and PC18K9) also agreed with major cell lineages and produced distinguishing marker genes with more rare types or subtypes reported (Figure 2C; Supplementary Figure 4). Finally, IKAP consumed less time (1hr 10m) than computing the 20 trial sets (5hr 13m) (Figure 2C). Although IKAP required an extra step to explore parameter space (19m), much time was saved because of fewer runs (3 candidate sets versus 20 trial sets) of time-consuming marker gene identification. The result shows that IKAP could help biological interpretation by picking appropriate parameters and reporting major cell groups that produce distinguishing marker genes within a reasonable time frame.

**Figure 2.**
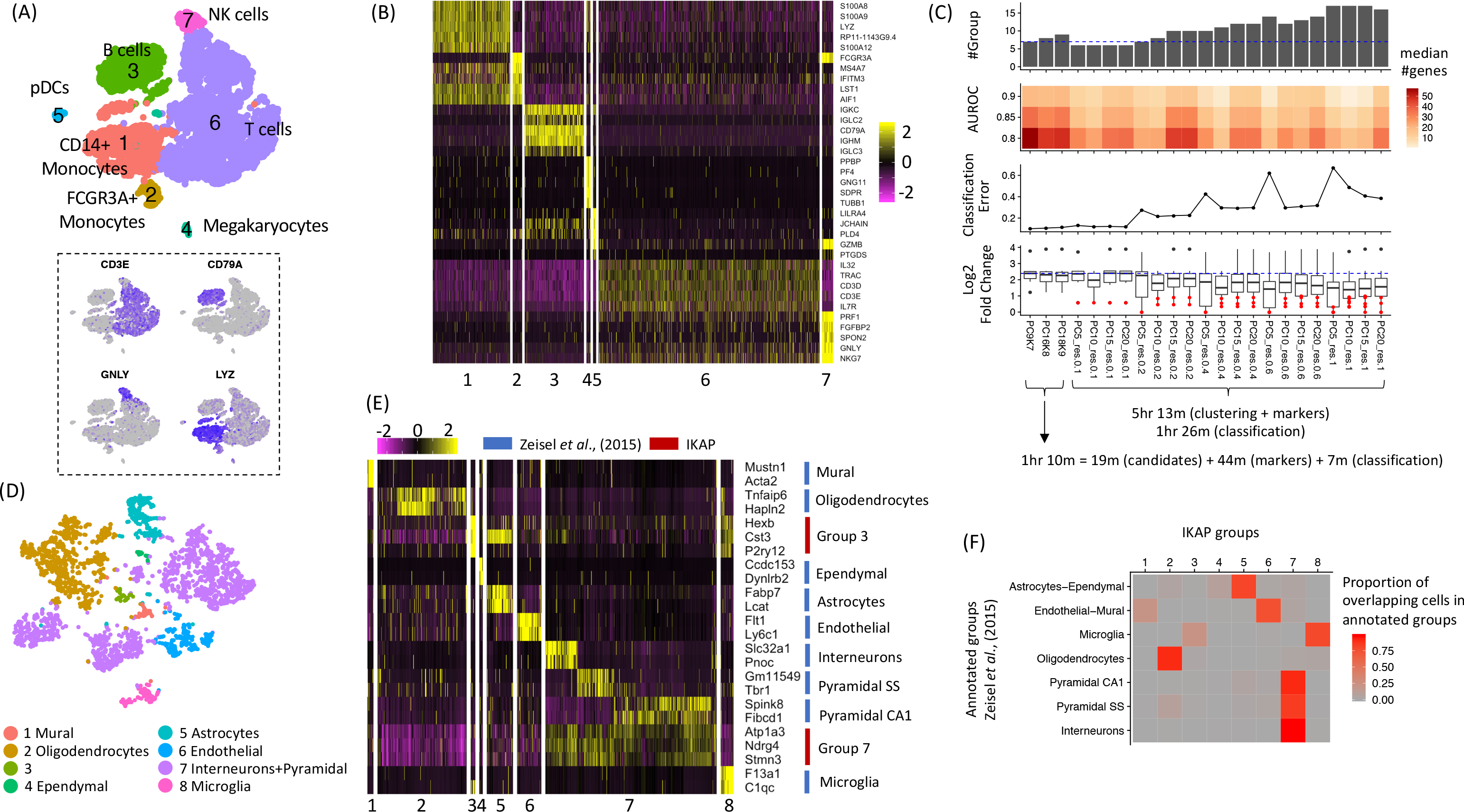
Major cell groups identified for PBMC_8K (A, B, and C) and the mouse cortex dataset (D, E, and F) were consistent with major lineages in PBMC and previously annotated cortex cell types respectively and well differentiated by marker genes. (A) Shown are tSNE plots for the 7 major groups identified by IKAP with cell lineages labeled (*top*) and expression of known marker genes (*bottom): CD3E* for T cells, *CD79A* for B cells, *GNLY* for NK cells, and *LYZ* for monocytes. (B) The heatmap for expression of the top 5 marker genes (by expression fold change) from each group in (A). Rows are genes and columns are cells. (C) Performance summary of 3 candidate sets proposed by IKAP and the 20 trial sets. Running time is shown at the bottom. (D) The tSNE plot for 8 major groups identified by IKAP in the mouse cortex dataset. (E) The heatmap for expression of marker genes annotated for major cell types in Zeisel *et al.*, 2015 (6) (*blue*) and marker genes identified by IKAP for groups 3 and 7 in (D) (*red*). (F) The heatmap indicates the proportion of overlapping cells between IKAP-identified major groups and major cell types annotated in Zeisel *et al.*, 2015 (6).

**Figure 3.**
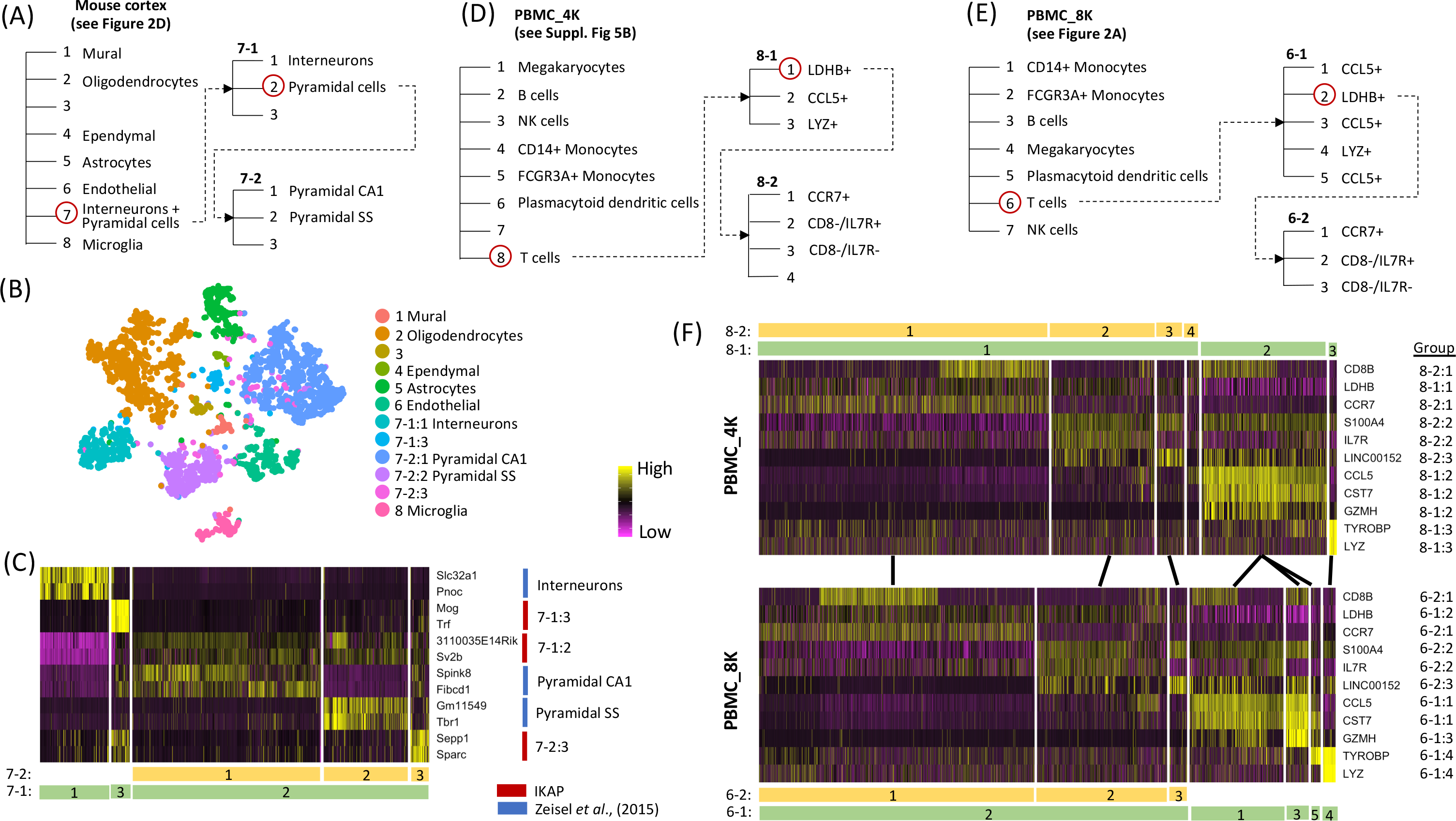
Examples of cell ontology proposed by IKAP. Three cell ontology examples were built by recursively running IKAP on the biggest groups (circled in red) for the mouse cortex dataset (A), PBMC_4K (D), and PBMC_8K (E). Putative cell types were labeled. Unknown types were left as blanks. (B) Shown is the tSNE plot for major groups and subgroups of group 7 presented in the mouse cortex ontology in (A). (C) The heatmap shows expression of marker genes identified by IKAP (*red*) and annotated in Zeisel *et al*., 2015 (6) (*blue*) for subgroups of group 7 in (A) (labeled at bottom). (F) Heatmaps show expression of selected marker genes that differentiate T cell subtypes in PBMC_4K (top; subgroups labeled according to the ontology in (D)) and in PBMC_8K (*bottom;* subgroups labeled according to the ontology in (E)). Subgroups with similar expression profiles are linked by lines between PBMC_4K and PBMC_8K.

To test robustness, we repeated the same analysis for another dataset PBMC_4K (~ 4K cells) from the same donor (5). This time two candidate sets (PC8K7 and PC20K8) were picked with PC20K8 marked as the best. Compared to the 20 trial sets, IKAP candidate sets were in better agreement with known cell lineages, known cell lineage marker genes being prioritized at the top, producing more distinguishing marker genes, and consuming shorter running time (Supplementary Figures 5-7).

Next we applied IKAP on a mouse cortex dataset (~ 3K cells) in which nine major cell types were previously annotated in Zeisel *et al.*, 2015 (6) (Supplementary Figure 8A). IKAP identified one candidate set with 8 major groups (PC13K8) (Figure 2D). Six groups were consistent with the annotated cell types as evidenced by expressing marker genes annotated specific to those cell types (Figure 2E) and high proportion of overlapping cells (Figure 2F). For the 2 remaining groups, one was a subtype of microglia cells (group 3 in Figure 2D) characterized by *Hexb* (AUROC=1.0), *Cst3* (0.98), and *P2ry12* (0.98) and the other was the union of 3 annotated cell types, interneurons, pyramidal SS and pyramidal CA1 (group 7 in Figure 2D), characterized by *Atp1a3* (0.93), *Ndrg4* (0.94), and *Stmn3* (0.93) (Figure 2E). Compared with the 20 trial sets (generated in the same way described above), marker genes in PC13K8 yielded higher expression fold change and lower classification error (Supplementary Figure 8B). Interestingly, the number of marker genes with high AUROC remained high as more subtypes were identified (which was not true for PBMC datasets), suggesting transcriptomic differentiation in the mouse cortex cells was very fine-grained. Overall, IKAP successfully recovered major cell types (rather than many subtypes) that were consistent with previous annotations and produced distinguishing marker genes.

Finally, a potential application of IKAP is to help redefine multi-layered cell ontology based on single-cell transcriptomics (7). IKAP can define one layer by splitting a cell population into *k* groups. By recursively applying IKAP within each group in the upper layer, subgroups can be identified in deeper layers. To demonstrate this approach, we expanded two more layers for all 3 datasets by applying IKAP on their biggest major groups and the biggest resulting subgroups. For the mouse cortex dataset, IKAP successfully recovered interneurons, pyramidal SS, and pyramidal CA1 by subdividing their union group (group 7 in Figure 2D) (Figure 3A-C). This 3-layer ontology (Figure 3A) offered a more complete view by not only differentiating all previously annotated cell types but also uncovering expression profiles shared across interneurons, pyramidal SS, and pyramidal CA1 such as expression of Atp1a3 across all three cell types (Figure 2E) and Sv2b across pyramidal cells (Figure 3C). For PBMC datasets, T cells (the biggest group) were subdivided into 2 layers of subgroups in which subgroups in the same layers were consistent between PBMC_4K and PBMC_8K, suggesting the ontology built by IKAP could be reproduced in replicates (Figure 3D-F; Supplementary Figures 9-10). Based on identified marker genes, subgroups in the 1^st^ layer were LDHB+, CCL5+, and LYZ+ T cells. The LDHB+ T cells were further divided into CCR7+, CD8-/IL7R+, and CD8-/IL7R-T cells in the 2^nd^ layer. Results shown above were the best sets selected among candidate sets proposed by IKAP. Ontology can be modified by manual inspection of different candidate sets. For example, within T cells, a more flattened ontology was achieved by using the candidate set with 7 subgroups (among 4 candidate sets) for PBMC_4K and the set with 10 subgroups (among 5 sets) for PBMC_8K (Supplementary Figures 11-13). In summary, although manual review may be still needed, IKAP could automate cell ontology curation using scRNA data by guiding multi-layered representations of cell groups – a task which has not been effectively addressed yet.

## Methods

### IKAP details

IKAP was implemented on top of Seurat (version 2.3.4) (3) in R (version 3.4). When running IKAP by default, it takes only a Seurat object which contains a normalized expression matrix and pre-computed covariates that need to be regressed out. The expression matrix will be scaled with covariates (if provided) regressed out using ScaleData function in Seurat. Then, IKAP finds variable genes using FindVariableGenes function in Seurat. All Seurat functions are run by default unless particular setting is specified. Default parameters for IKAP can be easily adjusted by users. Details are discussed in the following.

1. *Determine nPC*_*min*_, *nPC*_*maX*_, and *k*_*max*_. IKAP avoids specifying a particular number of top principal components (nPC) and *k* by exploring combinations of nPC and *k* (nPC, *k*). nPC_min_, nPC_max_, and *k*_*max*_ are used to define the search space of (nPC, *k*) such that the combinations (nPC*, *k\**) that can generate major groups are enclosed (*i.e*. nPC_min_ ≤ nPC* ≤ nPC_max_ and *k*\* ≤ *k*_*max*_). Setting large nPC_max_ and *k*_*max*_ increases the search space and, thus, the computation time. By following the concept of elbow method, nPC_min_ is computed as the first principal component (PC) such that decrease in explained standard deviation relative to the next PCs is less than 10% for all following PCs. By doing so, the top nPC_min_ PCs should contain informative features to define at least one set of major groups. Setting a nPC_max_ > nPC_min_ is for exploring more possible (nPC*, *k*\*) but would not affect the main result much. By default, nPC_max_ is set to nPC_min_ + 20. To set a *k*_*max*_ ≥ *k*\*, we found setting *r* > 1 in Seurat clustering usually produced many fine groups so by default, *k*_*max*_ is set to the average of the number of resulting groups using the top nPC_min_ PCs and that using the top nPC_max_ PCs by setting *r*_*ini*_ = 1.5. We varied the difference between nPC_max_ and nPC_min_ (nPC_max_ - nPC_min_ =10, 15, 20, and 25) and *r*_*ini*_ (=0.9, 1.2, 1.5, and 1.8) to generate 16 test sets for PBMC_4K, PBMC_8K, and the mouse cortex datasets and found grouping in the reported best sets did not change much (Supplementary Figures 14-16). This shows that our results were not sensitive to the default values of nPC_max_ and *r*_*ini*_.
2. *Generate k*_*max*_ *sets of groups for each nPC*. IKAP initializes the set of *k*_*ini*_ groups by setting r=1.0 using Seurat clustering. If *k*_*ini*_ < *k*_*max*_, increment *r* by 0.2 until *k*_*ini*_ >= *k*_*max*_. Two nearest groups measured by their centers in the PC space are merged iteratively, generating *k*_*ini*_ sets of groups but only the first *k*_*max*_ sets (with *k*=1 to *k*_*max*_) are used further.
3. *Compute gap statistic*. The gap statistic for a set of *k* groups is the difference between the log of sum of within-group pairwise distances over all *k* groups using the actual data and the log of expected sum of within-group pairwise distances over all *k* groups assuming data points (cells) are uniformly distributed in a bounded PC space where boundaries in each dimension are the minimum and the maximum of the actual data in that dimension. Details about gap statistic are described in (4).
4. *Compute marker genes*. IKAP utilizes FindAllMarker function in Seurat to compute marker genes. Only upregulated genes are reported. Other parameters are set by default.
5. *Build decision trees*. The idea of building the decision tree is to evaluate if a group of cells can be differentiated by considering multiple genes jointly. For each set of cell groups proposed by IKAP, a binary classifier (a decision tree) is built for each group using marker genes from all groups. The decision tree is built by R package rpart using default parameters.
6. *Compute classification error*. The R package rpart builds the decision tree for each group in a candidate set (see *Build decision trees* above) and also reports relative errors (*i.e*. training errors) at different number of splits (*nsplit*) along the decision tree. For each group at a given *nsplit*, the group-level classification error is computed as the product of the relative error and the fraction of cells in that group. The set-level classification error of a candidate set at a given *nsplit* is defined as the sum of all group-level classification errors. IKAP computes the final classification error for each candidate set as the average of set-level classification errors for *nsplit* = 5 to 15. Note that the tree usually did not grow more than 15 splits in the experiments shown in this study.
7. *Select the best and alternative candidate sets*. The workflow of selecting candidate sets (PC9K7, PC16K8, and PC18K9) for PBMC_8K is shown in Supplementary Figure 17. The formal procedure is briefly described as follows. By computing gap increase from a set of *k-1* groups to *k* groups (see Step 3 in Figure 1) for every tested nPC, IKAP generates a gap-increase matrix *M* in which rows correspond to nPC and columns correspond to *k*. Note that each combination of (nPC, *k*) corresponds to a set of cell groups. IKAP first filters out those (nPC, *k)’s* with gap increase ≤ mean + standard deviation. Then, IKAP picks the largest non-zero gap increase for every *k* (every column of M), generating a list of gap increases and a list of corresponding (nPC, *k)’s* where *k’s* are different. The list of (nPC, *k)’s* is sorted by corresponding gap increases in descending order. The first (nPC, *k*) (which corresponds to the largest gap increase) is picked as a candidate set. Then, IKAP goes down to the list one by one and adds the (nPC, *k*) to the candidate list if the nPC and *k* are greater than all nPCs and *k’s* already in the candidate list. This requirement is to look for cases where additional cell groups (larger *k*) are identified because of incorporating additional PCs (larger nPC). Finally, among selected candidate sets, the one with the lowest classification error (see *Compute classification error*) is marked as the best and the rest are alternatives.

### Performance summary

The AUROC (Area Under the ROC curve) was computed using R package PRROC. We counted the median of numbers of genes with high AUROC (> 0.8, 0.85, and 0.9) across all groups. The classification error was computed as described above (*Compute classification error* in **IKAP details)**. Average expression log fold change (AvgLFC) was reported by Seurat for each marker gene in each group. For each group, we sorted genes by AvgLFC and only considered marker genes with AUROC > 0.8. Among those, we computed the mean of AvgLFC across top 10 (or n if n < 10) marker genes for each group. Running time was measured on 4.2 GHz Intel Core i7 iMac desktop with 64 GB memory.

### PBMC_4K and PBMC_8K datasets

PBMC_4K and PBMC_8K were downloaded from the 10x Genomics website (5). They were filtered and normalized using the R package Seurat (3). We removed cells with less than 200 genes expressed or the unique molecular identifier (UMI) count of mitochondrial genes > 5% of the total UMI count. For each dataset, we regressed out the percentage of mitochondrial gene UMI count and the total UMI count from the normalized expression matrix and scaled the matrix using ScaleData function in Seurat. Finally, we got the expression matrix with 16,746 genes and 4,077 cells for PBMC_4K and the matrix with 18,408 genes and 8090 cells for PBMC_8K.

### Mouse cortex dataset

The dataset was obtained from (6). We normalized and scaled the expression matrix as we did for PBMC_4K and PBMC_8K but we did not filter out any cells in order to be consistent with the published work. In total, the expression matrix comprised 19,972 genes and 3,005 cells.

### Cell lineage recognition

Major cell groups in PBMC datasets (Figure 2A and Figure 3D-E) were annotated based on expression of marker genes and literature. CD14+ monocytes: expression of *LYZ* and *S100A8* (8). FCGR3A+ monocytes: expression of *FCGR3A* and *MS4A7*, a monocyte lineage marker (9). B cells: expression of *CD79A* (10). Megakaryocytes: expression of PPBP (11). Plasmacytoid dendritic cells: expression of *LILRA4* (12). T cells: expression of *CD3E* (13). NK cells: expression of *GNLY* but not *CD3E* (14).

### Data availability

PBMC_4K and PBMC_8K were downloaded from 10x Genomics website: https://www.10xgenomics.com/resources/datasets/. The mouse cortex dataset was acquired from the accession numbers provided in the original publications.

### Software availability

The R script of IKAP is available at: https://github.com/NHLBI-BCB/IKAP

## Author Contributions

M.P. and Y.C. conceived the study. Y.C. developed and implemented the algorithm and drafted the manuscript; A.S. and F.S. helped with implementation; C.U., C.S., K.S., and A.W. helped with cell type annotation. M.P. supervised the research. All authors reviewed and approved the manuscript.

## Supplementary figure legends

**Supplementary Figure 1. Gap statistics increased as more cell groups were identified in PBMC_8K using different numbers of top principal components (nPCs) with large gap increase seen around the number of groups (k) = 7, 8, or 9**.

**Supplementary Figure 2. Major cell lineages in PBMC were not well aligned with the 20 trial sets of cell groups generated for PBMC_8K by varying resolution (*r*) and the number of top principal components (nPC) using Seurat clustering**. (A) The tSNE plots for the 20 trial sets. (B) Expression of PBMC lineage marker genes: *CD3E* for T cells, *CD79A* for B cells, GNLY for NK cells and LYZ for monocytes.

**Supplementary Figure 3. Marker gene expression for 6 trial sets selected from the 20 trial sets for PBMC_8K shown in Supplementary Figure 2**. In the heatmaps, rows are genes and columns are cells. Groups of cells (separated by vertical white lines) from left to right correspond to groups of corresponding trial sets in Supplementary Figure 2 in the order of 0, 1, 2, … etc.

**Supplementary Figure 4. Two alternative sets (PC16K8 and PC18K9) of major cell groups identified for PBMC_8K by IKAP**. Shown are tSNE plots of the major groups (*left*) and heatmaps for expression of top marker genes (by expression fold change) (*right*) for PC16K8 (*top*) and PC18K9 (*bottom*). Rows are genes and columns are cells in the heatmaps.

**Supplementary Figure 5. Two candidate sets (PC8K7 and PC20K8) of major groups identified for PBMC_4K by IKAP were aligned with major cell lineages in PBMC and well differentiated by lineage marker genes**. tSNE plots show major groups of PC8K7 (A) and PC20K8 (B). Heatmaps show expression of top marker genes (by expression fold change) of each group for PC8K7 (C) and PC20K8 (D). (E) tSNE plots for expression of lineage marker genes: *CD3E* for T cells, *CD79A* for B cells, *GNLY* for NK cells and *LYZ* for monocytes. (F) Performance summary of the 2 candidate sets proposed by IKAP and 20 trial sets. Running time is shown at the bottom.

**Supplementary Figure 6. Major cell lineages in PBMC were not well aligned with the 20 trial sets of cell groups generated for PBMC_4K by varying resolution (*r*) and the number of top principal components (nPC) using Seurat clustering**. (*A) The tSNE plots for the 20 trial sets. (B) Expression of PBMC cell lineage marker genes: CD3E for T cells, CD79A for B cells, GNLY for NK cells and LYZ for monocytes*.

**Supplementary Figure 7. Expression of top marker genes for 6 trial sets selected from the 20 trial sets for PBMC_4K shown in Supplementary Figure 6**. In the heatmaps, rows are genes and columns are cells. Groups of cells (separated by vertical white lines) from left to right correspond to groups of corresponding trial sets in Supplementary Figure 6 in the order of 0, 1, 2, … etc.

**Supplementary Figure 8. Comparison among previously annotated major cell types, major cell groups identified by IKAP, and the 20 trial sets of cell groups for the mouse cortex data**. (*A*) The top tSNE plot shows 9 cell types with original labels in Zeisel *et al*., 2015 (6) where astrocytes were merged with ependymal and endothelial merged with mural. The modified tSNE plot at bottom recovered ependymal and mural types using group 4 and group 1 identified by IKAP in Figure 2D. (B) Performance summary of the set proposed by IKAP, the 20 trial sets, and the modified version of major cell types in (A). Running time is shown at the bottom.

**Supplementary Figure 9. An example of 2-layer T cell ontology proposed by IKAP for PBMC_4K**. Shown on the left is the ontology with two layers (also shown in Figure 3D). The heatmap shows expression of top marker genes (ranked by expression fold change) of each subgroup. Rows are genes and columns are cells.

**Supplementary Figure 10. An example of 2-layer T cell ontology proposed by IKAP for PBMC_8K**. Shown on the left is the ontology with two layers (also shown in Figure 3E). The heatmap shows expression of top marker genes (ranked by expression fold change) of each subgroup. Rows are genes and columns are cells.

**Supplementary Figure 11. Alternative sets of T cell subgroups reported by IKAP for PBMC_4K and PBMC_8K**. Two PBMC ontologies with T cell subgroups are shown on the left for PBMC_4K (*top*) and PBMC_8K (*bottom*). Expression of marker genes is plotted in heatmaps on the right. Rows are genes and columns are cells. Subgroups with similar expression profiles are linked by lines.

**Supplementary Figure 12. Expression of top marker genes (ranked by expression fold change) for PBMC_4K T cell subgroups shown in Supplementary Figure 11**. *Rows are genes and columns are cells*.

**Supplementary Figure 13. Expression of top marker genes (ranked by expression fold change) for PBMC_8K T cell subgroups shown in Supplementary Figure 11**. *Rows are genes and columns are cells*.

**Supplementary Figure 14. Major cell groups identified by IKAP were not sensitive to settings of parameters, r_ini_ and (nPC_max_-nPC_min_), for PBMC_4K (see Online Methods)**. *The tSNE plots for 16 sets of major cell groups generated by varying r_ini_ (=0.9, 1.2, 1.5, and 1.8) and (nPC_max_-nPC_min_) (=10, 15, 20, and 25) using PBMC_4K. By default r_ini_=1.5 and (nPC_max_-nPC_min_)=20*.

**Supplementary Figure 15. Major cell groups identified by IKAP were not sensitive to settings of parameters, *r*_*ini*_ and (nPC_max_-nPC_min_), for PBMC_8K (see Online Methods)**. *The tSNE plots for 16 sets of major cell groups generated by varying r_ini_ (=0.9, 1.2, 1.5, and 1.8) and (nPC_max_-nPC_min_) (=10, 15, 20, and 25) using PBMC_8K. By default r_ini_=1.5 and (nPC_max_-nPC_min_)=20*.

**Supplementary Figure 16. Major cell groups identified by IKAP were not sensitive to settings of parameters, *r*_*ini*_ and (*nPC_max_-nPC_min_*), for the mouse cortex dataset (see Online Methods).** The tSNE plots for 16 sets of major cell groups generated by varying *r*_*ini*_ (=0.9, 1.2, 1.5, and 1.8) and (nPC_max_-nPC_min_) (=10, 15, 20, and 25) using the mouse cortex dataset. By default *r*_*ini*_=1.5 and (nPC_max_-nPC_min_)=20.

**Supplementary Figure 17. The workflow of selecting candidate sets (PC9K7, PC16K8, and PC18K9) for PBMC_8K**. Given a gap-increase matrix *M* (see Figure 1 for how to compute gap increase), the following steps were taken. Step **0:** filter entries by gap increases > mean + standard deviation. Step 1: take the max gap increase across rows for each column (*k*) and record the corresponding (nPC, *k*). Step **2:** sort recorded (nPC, *k)’s* based on their corresponding gap increases. Step 3: add the first (nPC, *k*), which is PC9K7, into the candidate list. Step 4: remove the second (nPC, *k*), which is PC9K6, because its nPC (=9) is not larger than nPC of the candidate (=9) in the candidate list and neither is its *k* not larger than *k* of the candidate (=7) in the candidate list. Step 5: add the third (nPC, *k*) into the candidate list because its nPC (=16) is larger than nPC of the candidate (=9) in the candidate list and so is its *k*. Step 6: add the fourth (nPC, *k*) into the candidate list because its nPC **(=20)** is larger than all nPCs of the candidates (=9 and 16) in the candidate list and so is its *k*. Finally, PC9K7, PC16K8, and PC18K9 were selected as candidate sets for PBMC_8K.

## References

1. Andrews TS, Hemberg M. Identifying cell populations with scRNASeq. Mol Aspects Med. 2018;59:114–22. doi:10.1016/j.mam.2017.07.002

2. Kiselev VY, Andrews TS, Hemberg M. Challenges in unsupervised clustering of single-cell RNA-seq data. Nat Rev Genet. 2019. doi:10.1038/s41576-018-0088-9

3. Butler A, Hoffman P, Smibert P, Papalexi E, Satija R. Integrating single-cell transcriptomic data across different conditions, technologies, and species. Nat Biotechnol. 2018;36(5):411–20. doi:10.1038/nbt.4096

4. Tibshirani R, Walther G, Hastie T. Estimating the number of clusters in a data set via the gap statistic. J Roy Stat Soc B. 2001;63:411–23. doi:Doi 10.1111/1467-9868.00293

5. Genomics x. Support: single cell gene expression datasets. 10x Genomics. 2017.

6. Zeisel A, Munoz-Manchado AB, Codeluppi S, Lonnerberg P, La Manno G, Jureus A, et al. Brain structure. Cell types in the mouse cortex and hippocampus revealed by single-cell RNA-seq. Science. 2015;347(6226):1138–42. doi:10.1126/science.aaa1934

7. Bakken T, Cowell L, Aevermann BD, Novotny M, Hodge R, Miller JA, et al. Cell type discovery and representation in the era of high-content single cell phenotyping. BMC Bioinformatics. 2017;18(Suppl 17):559. doi:10.1186/s12859-017-1977-1

8. Zawada AM, Rogacev KS, Rotter B, Winter P, Marell RR, Fliser D, et al. SuperSAGE evidence for CD14++CD16+ monocytes as a third monocyte subset. Blood. 2011;118(12):e50–61. doi:10.1182/blood-2011-01-326827

9. Gingras MC, Lapillonne H, Margolin JF. CFFM4: a new member of the CD20/FcepsilonRIbeta family. Immunogenetics. 2001;53(6):468–76. doi:10.1007/s002510100345

10. Chu PG, Arber DA. CD79: a review. Appl Immunohistochem Mol Morphol. 2001;9(2):97–106.

11. Zhang C, Gadue P, Scott E, Atchison M, Poncz M. Activation of the megakaryocyte-specific gene platelet basic protein (PBP) by the Ets family factor PU.1. J Biol Chem. 1997;272(42):26236–46.

12. Cho M, Ishida K, Chen J, Ohkawa J, Chen W, Namiki S, et al. SAGE library screening reveals ILT7 as a specific plasmacytoid dendritic cell marker that regulates type I IFN production. Int Immunol. 2008;20(1):155–64. doi:10.1093/intimm/dxm127

13. Chetty R, Gatter K. CD3: structure, function, and role of immunostaining in clinical practice. J Pathol. 1994;173(4):303–7. doi:10.1002/path.1711730404

14. Pena SV, Krensky AM. Granulysin, a new human cytolytic granule-associated protein with possible involvement in cell-mediated cytotoxicity. Semin Immunol. 1997;9(2):117–25. doi:10.1006/smim.1997.0061

