## Supplementary Figures for "IKAP - Identifying K mAjor cell Population groups in single-cell RNA-seq analysis"

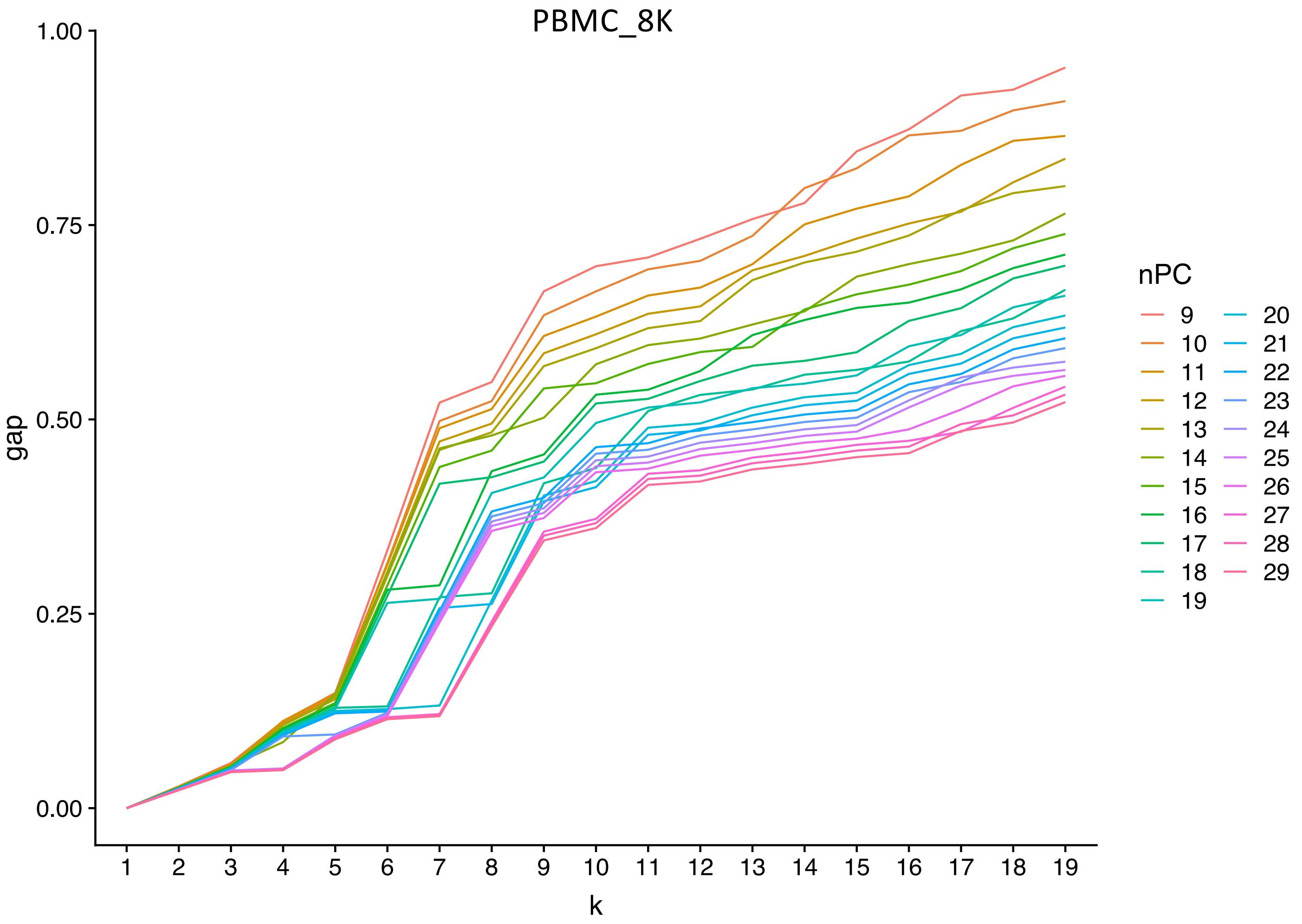

PBMC_8K
1.00
0.75
0.50
0.25
0.00
gap
1 2 3 4 5 6 7 8 9 10 11 12 13 14 15 16 17 18 19
k
nPC

(A)

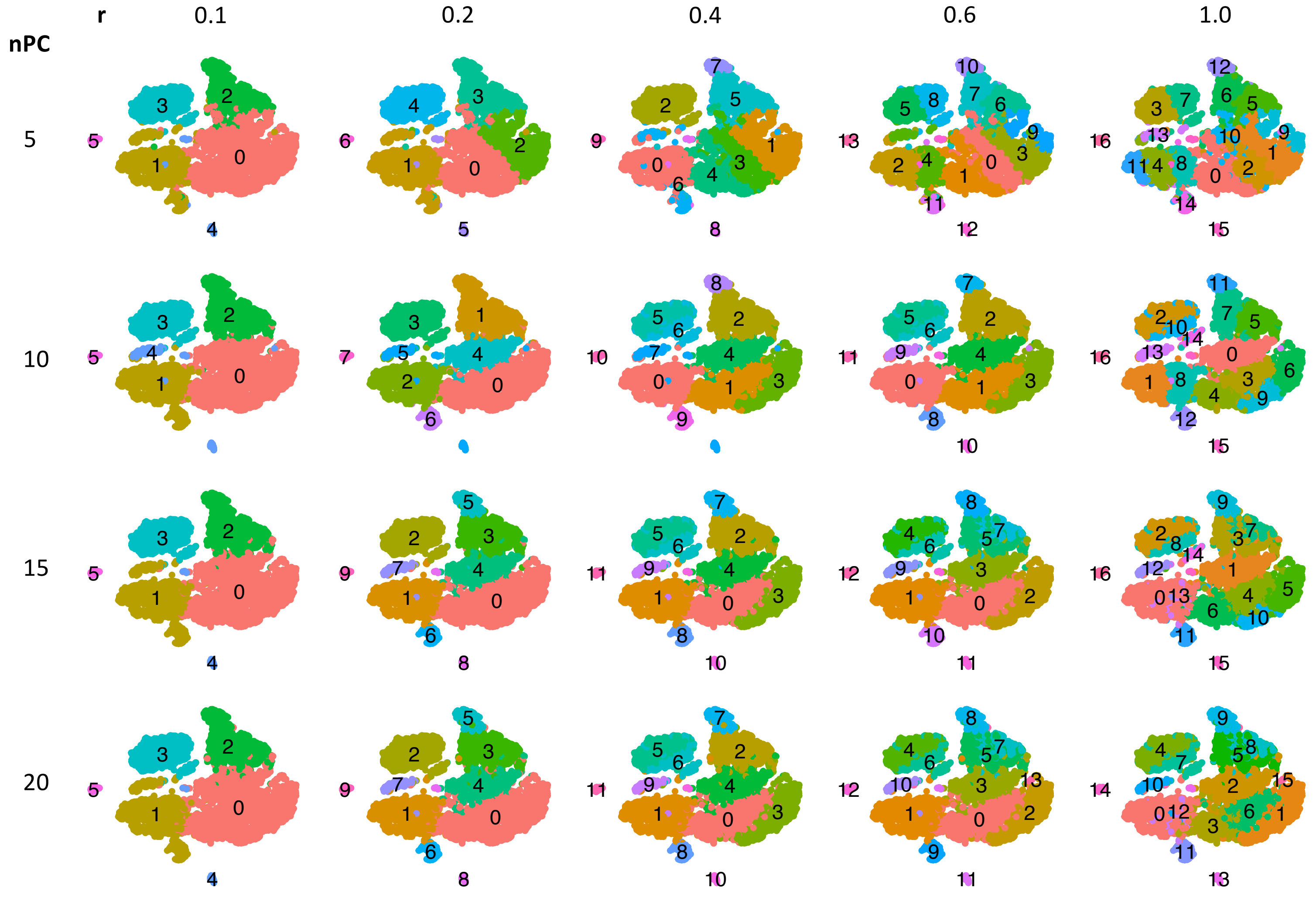

(B)

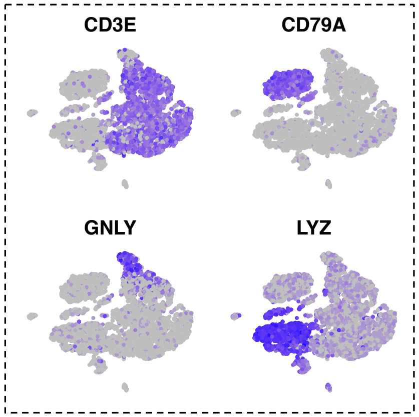

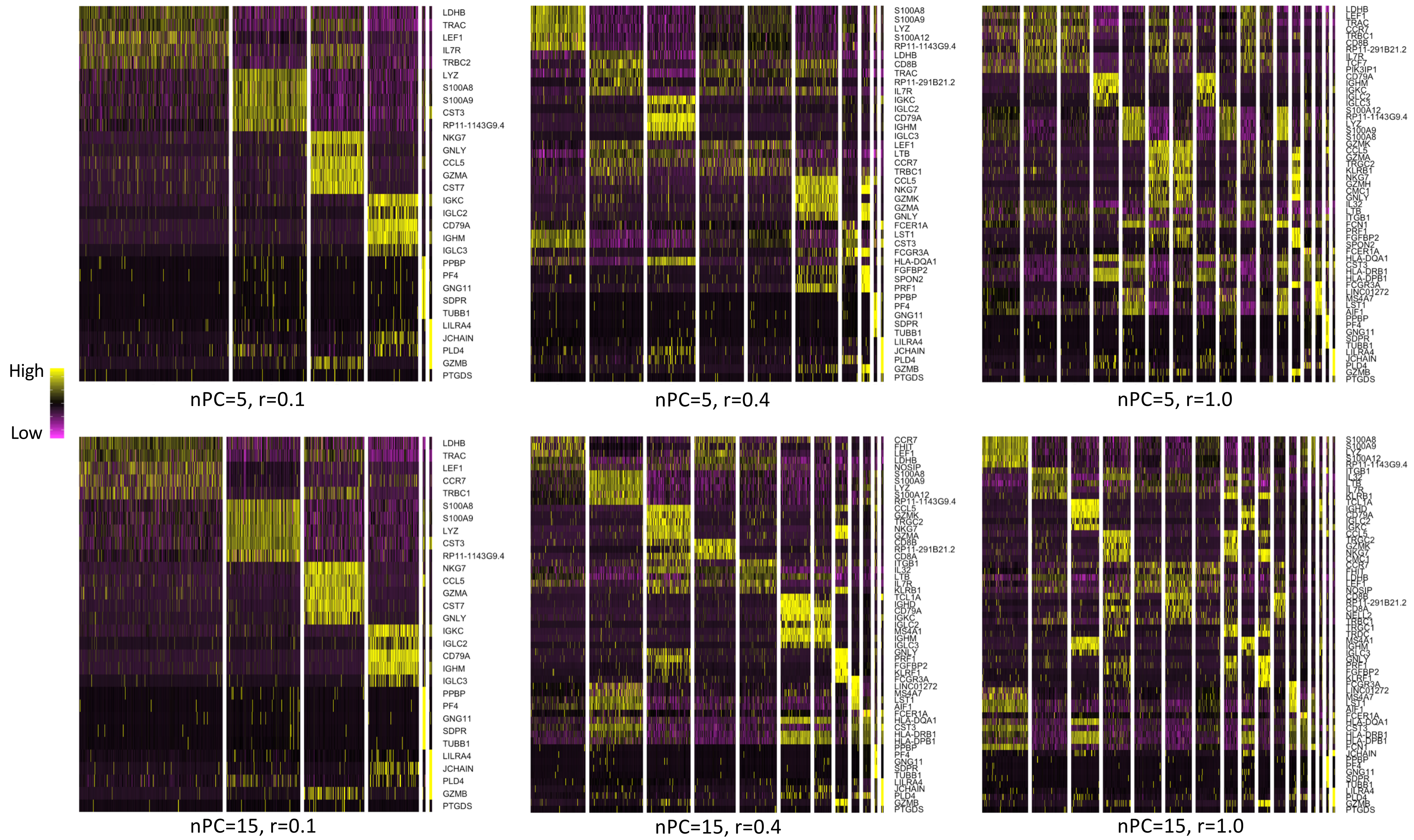
High
Low
nPC=5, r=0.1
LDHB
TRAC
LEF1
IL7R
TRBC2
LYZ
S100A8
S100A9
CST3
RP11-1143G9.4
NKG7
GNLY
CCL5
GZMA
CST7
IGKC
IGLC2
CD79A
IGHM
IGLC3
PPBP
PF4
GNG11
SDPR
TUBB1
LILRA4
JCHAIN
PLD4
GZMB
PTGDS
nPC=5, r=0.4
S100A8
S100A9
LYZ
S100A12
RP11-1143G9.4
LDHB
CD8B
TRAC
RP11-291B21.2
IL7R
IGKC
IGLC2
CD79A
IGHM
IGLC3
LEF1
LTB
CCR7
TRBC1
CCL5
NKG7
GZMK
GZMA
GNLY
FCER1A
LST1
CST3
FCGR3A
HLA-DQA1
FGFBP2
SPON2
PRF1
PPBP
PF4
GNG11
SDPR
TUBB1
LILRA4
JCHAIN
PLD4
GZMB
PTGDS
nPC=5, r=1.0
nPC=15, r=0.1
LDHB
TRAC
LEF1
CCR7
TRBC1
S100A8
S100A9
LYZ
CST3
RP11-1143G9.4
NKG7
CCL5
GZMA
CST7
GNLY
IGKC
IGLC2
CD79A
IGHM
IGLC3
PPBP
PF4
GNG11
SDPR
TUBB1
LILRA4
JCHAIN
PLD4
GZMB
PTGDS
nPC=15, r=0.4
nPC=15, r=1.0

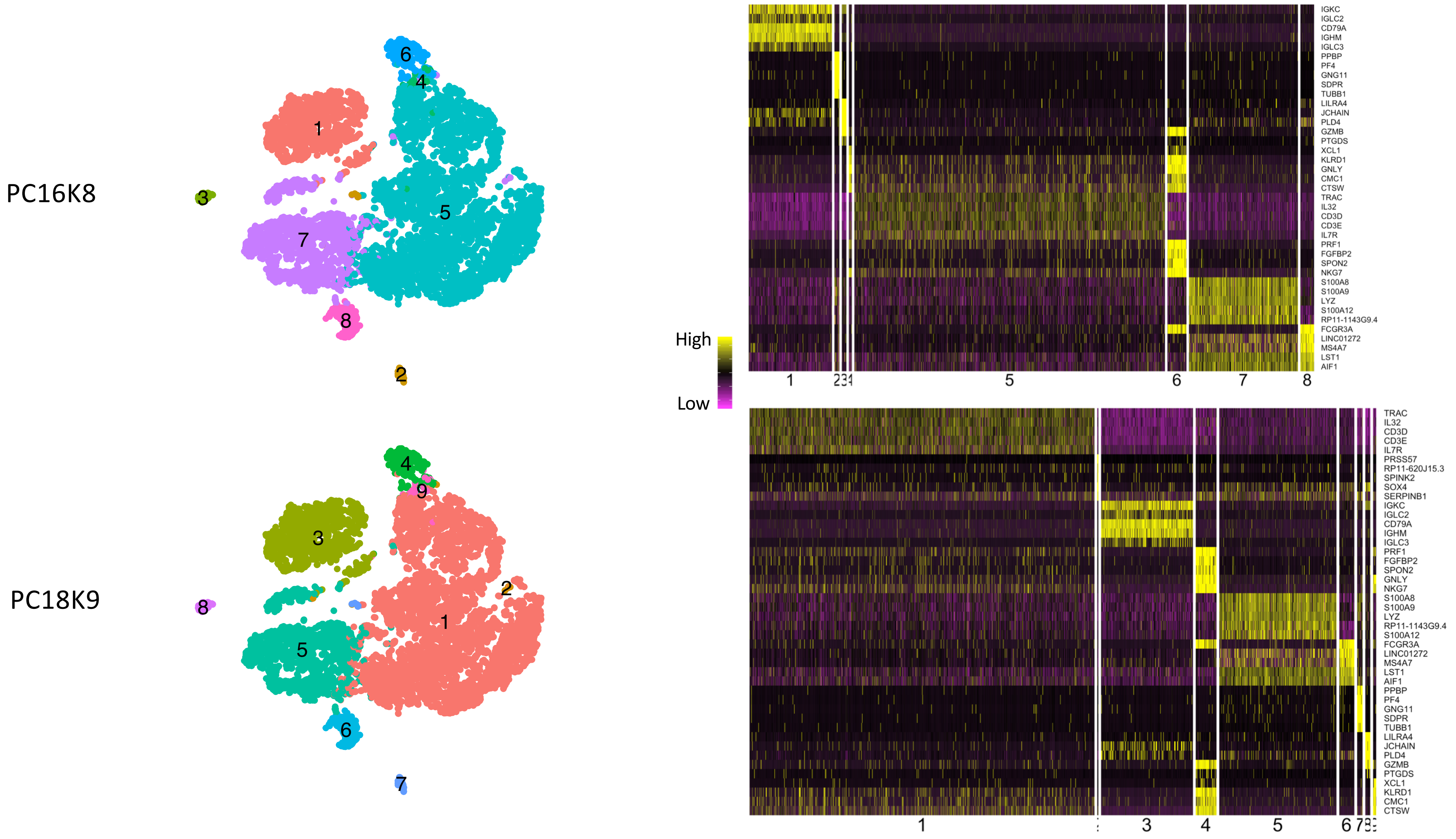
PC16K8
PC18K9
1
2
3
4
5
6
7
8
9
High
Low
IGKC
IGLC2
CD79A
IGHM
IGLC3
PPBP
PF4
GNG11
SDPR
TUBB1
LILRA4
JCHAIN
PLD4
GZMB
PTGDS
XCL1
KLRD1
GNLY
CMC1
CTSW
TRAC
IL32
CD3D
CD3E
IL7R
PRF1
FGFBP2
SPON2
NKG7
S100A8
S100A9
LYZ
S100A12
RP11-1143G9.4
FCGR3A
LINC01272
MS4A7
LST1
AIF1
TRAC
IL32
CD3D
CD3E
IL7R
PRSS57
RP11-620J15.3
SPINK2
SOX4
SERPINB1
IGKC
IGLC2
CD79A
IGHM
IGLC3
PRF1
FGFBP2
SPON2
GNLY
NKG7
S100A8
S100A9
LYZ
RP11-1143G9.4
S100A12
FCGR3A
LINC01272
MS4A7
LST1
AIF1
PPBP
PF4
GNG11
SDPR
TUBB1
LILRA4
JCHAIN
PLD4
GZMB
PTGDS
XCL1
KLRD1
CMC1
CTSW

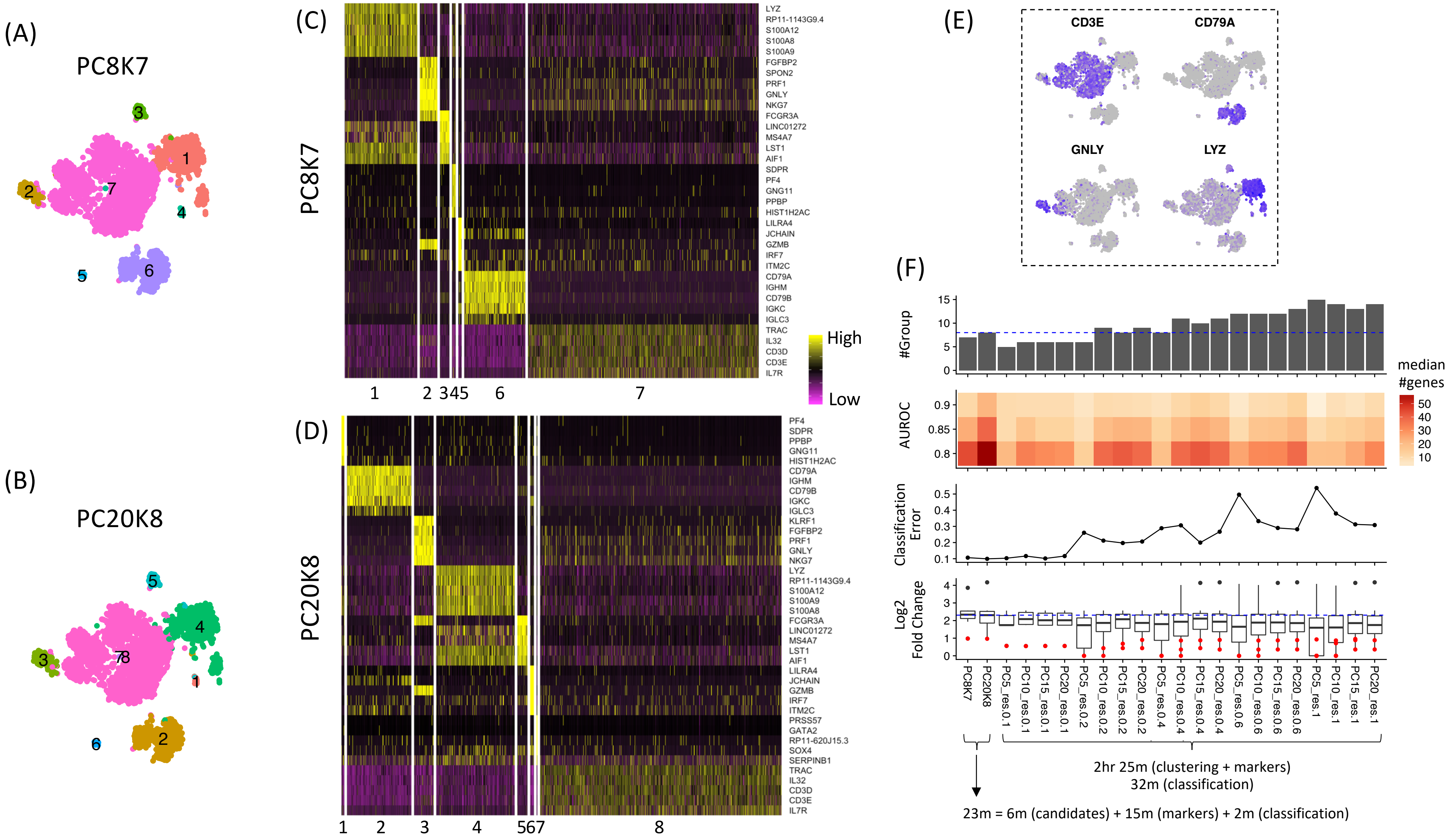

(A)
PC8K7
(B)
PC20K8
(C)
PC8K7
LYZ
RP11-1143G9.4
S100A12
S100A8
S100A9
FGFBP2
SPON2
PRF1
GNLY
NKG7
FCGR3A
LINC01272
MS4A7
LST1
AIF1
SDPR
PF4
GNG11
PPBP
HIST1H2AC
LILRA4
JCHAIN
GZMB
IRF7
ITM2C
CD79A
IGHM
CD79B
IGKC
IGLC3
TRAC
IL32
CD3D
CD3E
IL7R
1 2 345 6 7
High
Low
(D)
PC20K8
PF4
SDPR
PPBP
GNG11
HIST1H2AC
CD79A
IGHM
CD79B
IGKC
IGLC3
KLRF1
FGFBP2
PRF1
GNLY
NKG7
LYZ
RP11-1143G9.4
S100A12
S100A9
S100A8
FCGR3A
LINC01272
MS4A7
LST1
AIF1
LILRA4
JCHAIN
GZMB
IRF7
ITM2C
PRSS57
GATA2
RP11-620J15.3
SOX4
SERPINB1
TRAC
IL32
CD3D
CD3E
IL7R
1 2 3 4 567 8
(E)
CD3E
CD79A
GNLY
LYZ
(F)
#Group
AUROC
median #genes
50 40 30 20 10
Classification Error
Log2 Fold Change
PC8K7
PC20K8
PC5_res.0.1
PC10_res.0.1
PC15_res.0.1
PC20_res.0.1
PC5_res.0.2
PC10_res.0.2
PC15_res.0.2
PC20_res.0.2
PC5_res.0.4
PC10_res.0.4
PC15_res.0.4
PC20_res.0.4
PC5_res.0.6
PC10_res.0.6
PC15_res.0.6
PC20_res.0.6
PC5_res.1
PC10_res.1
PC15_res.1
PC20_res.1
2hr 25m (clustering + markers)
32m (classification)
23m = 6m (candidates) + 15m (markers) + 2m (classification)

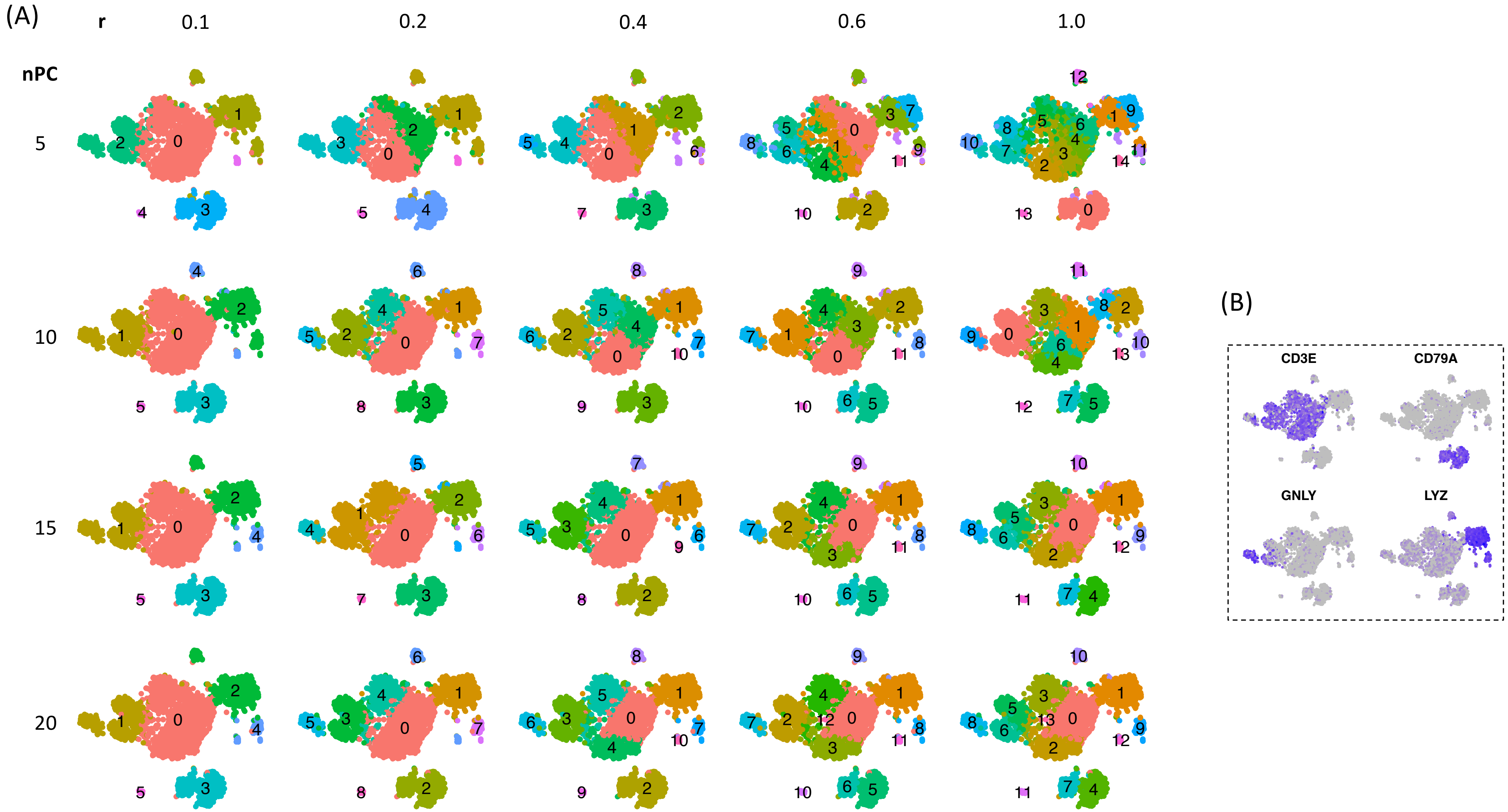
(A)
r
0.1
0.2
0.4
0.6
1.0
nPC
5
10
15
20
(B)
CD3E
CD79A
GNLY
LYZ

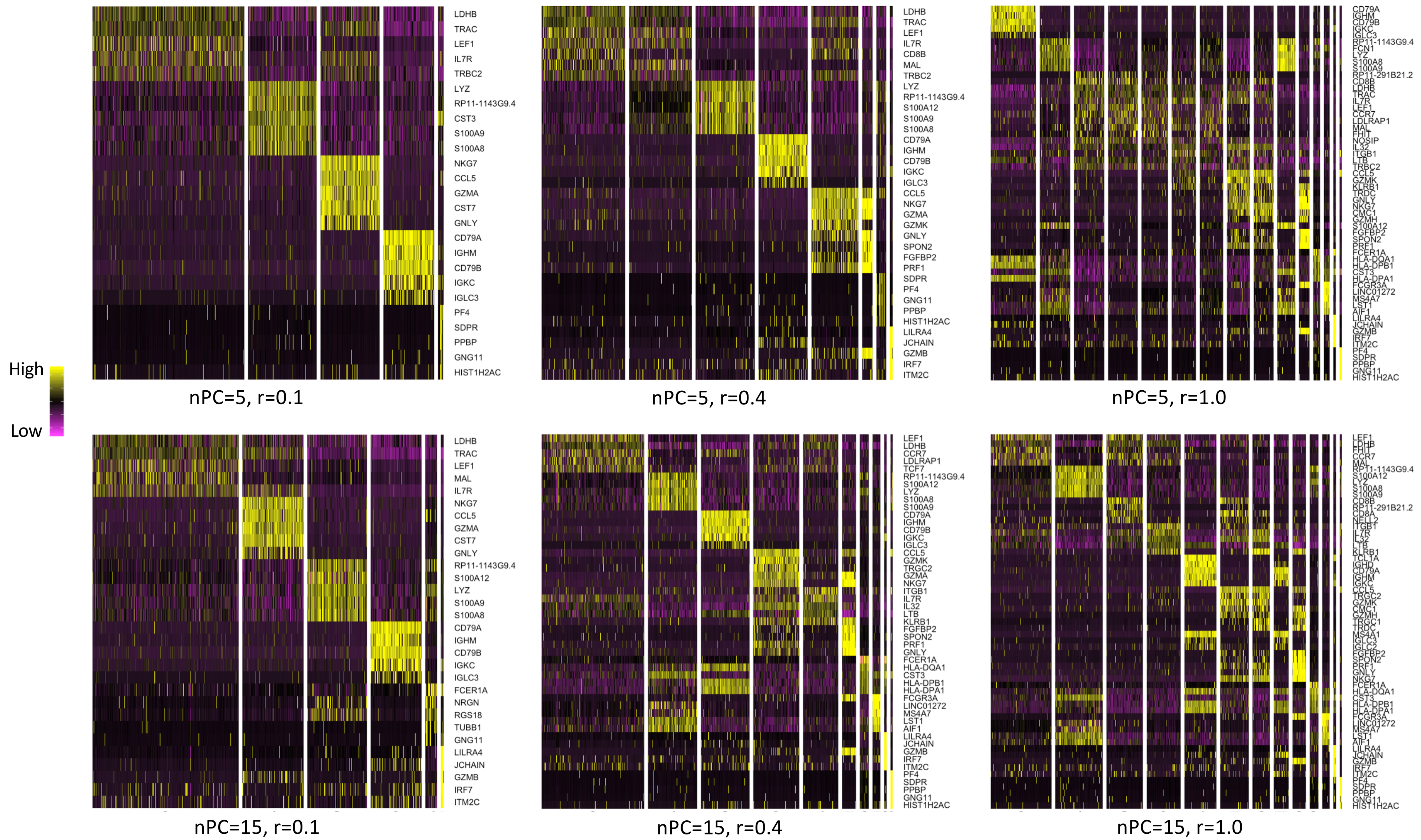

High
Low
LDHB
TRAC
LEF1
IL7R
TRBC2
LYZ
RP11-1143G9.4
CST3
S100A9
S100A8
NKG7
CCL5
GZMA
CST7
GNLY
CD79A
IGHM
CD79B
IGKC
IGLC3
PF4
SDPR
PPBP
GNG11
HIST1H2AC
nPC=5, r=0.1
LDHB
TRAC
LEF1
IL7R
CD8B
MAL
TRBC2
LYZ
RP11-1143G9.4
S100A12
S100A9
S100A8
CD79A
IGHM
CD79B
IGKC
IGLC3
CCL5
NKG7
GZMA
GZMK
GNLY
SPON2
FGFBP2
PRF1
SDPR
PF4
GNG11
PPBP
HIST1H2AC
LILRA4
JCHAIN
GZMB
IRF7
ITM2C
nPC=5, r=0.4
CD79A
IGHM
CD79B
IGKC
IGLC3
RP11-1143G9.4
FCN1
LYZ
S100A8
S100A9
RP11-291B21.2
CD8B
LDHB
TRAC
IL7R
LEF1
CCR7
LDLRAP1
MAL
FHIT
NOSIP
IL32
ITGB1
LTB
TRBC2
CCL5
GZMK
KLRB1
TRDC
GNLY
NKG7
CMC1
GZMH
S100A12
FGFBP2
SPON2
PRF1
FCER1A
HLA-DQA1
HLA-DPB1
CST3
HLA-DPA1
FCGR3A
LINC01272
MS4A7
LST1
AIF1
LILRA4
JCHAIN
GZMB
IRF7
ITM2C
PF4
SDPR
PPBP
GNG11
HIST1H2AC
nPC=5, r=1.0
LDHB
TRAC
LEF1
MAL
IL7R
NKG7
CCL5
GZMA
CST7
GNLY
RP11-1143G9.4
S100A12
LYZ
S100A9
S100A8
CD79A
IGHM
CD79B
IGKC
IGLC3
FCER1A
NRGN
RGS18
TUBB1
GNG11
LILRA4
JCHAIN
GZMB
IRF7
ITM2C
nPC=15, r=0.1
LEF1
LDHB
CCR7
LDLRAP1
TCF7
RP11-1143G9.4
S100A12
LYZ
S100A8
S100A9
CD79A
IGHM
CD79B
IGKC
IGLC3
CCL5
GZMK
TRGC2
GZMA
NKG7
ITGB1
IL7R
IL32
LTB
KLRB1
FGFBP2
SPON2
PRF1
GNLY
FCER1A
HLA-DQA1
CST3
HLA-DPB1
HLA-DPA1
FCGR3A
LINC01272
MS4A7
LST1
AIF1
LILRA4
JCHAIN
GZMB
IRF7
ITM2C
PF4
SDPR
PPBP
GNG11
HIST1H2AC
nPC=15, r=0.4
LEF1
LDHB
FHIT
CCR7
MAL
RP11-1143G9.4
S100A12
LYZ
S100A8
S100A9
CD8B
RP11-291B21.2
CD8A
NELL2
ITGB1
IL7R
IL32
LTB
KLRB1
TCL1A
IGHD
CD79A
IGHM
IGKC
CCL5
TRGC2
GZMK
CMC1
GZMH
TRGC1
TRDC
MS4A1
IGLC3
IGLC2
FGFBP2
SPON2
PRF1
GNLY
NKG7
FCER1A
HLA-DQA1
CST3
HLA-DPB1
HLA-DPA1
FCGR3A
LINC01272
MS4A7
LST1
AIF1
LILRA4
JCHAIN
GZMB
IRF7
ITM2C
PF4
SDPR
PPBP
GNG11
HIST1H2AC
nPC=15, r=1.0

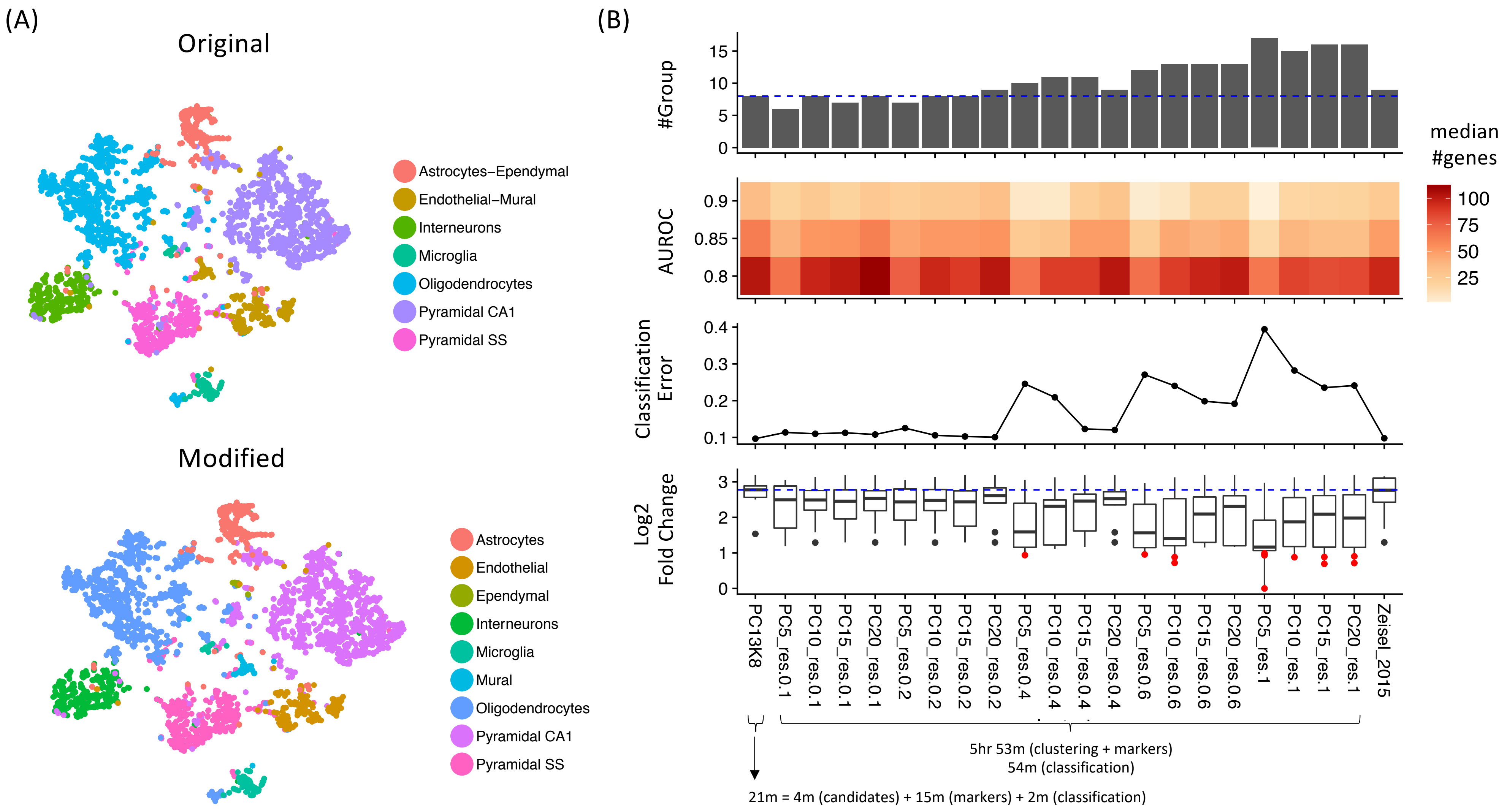

(A)
Original
Astrocytes–Ependymal
Endothelial–Mural
Interneurons
Microglia
Oligodendrocytes
Pyramidal CA1
Pyramidal SS
Modified
Astrocytes
Endothelial
Ependymal
Interneurons
Microglia
Mural
Oligodendrocytes
Pyramidal CA1
Pyramidal SS
(B)
#Group
AUROC
Classification Error
Log2 Fold Change
median #genes
PC13K8
PC5_res.0.1
PC10_res.0.1
PC15_res.0.1
PC20_res.0.1
PC5_res.0.2
PC10_res.0.2
PC15_res.0.2
PC20_res.0.2
PC5_res.0.4
PC10_res.0.4
PC15_res.0.4
PC20_res.0.4
PC5_res.0.6
PC10_res.0.6
PC15_res.0.6
PC20_res.0.6
PC5_res.1
PC10_res.1
PC15_res.1
PC20_res.1
Zeisel_2015
5hr 53m (clustering + markers)
54m (classification)
21m = 4m (candidates) + 15m (markers) + 2m (classification)

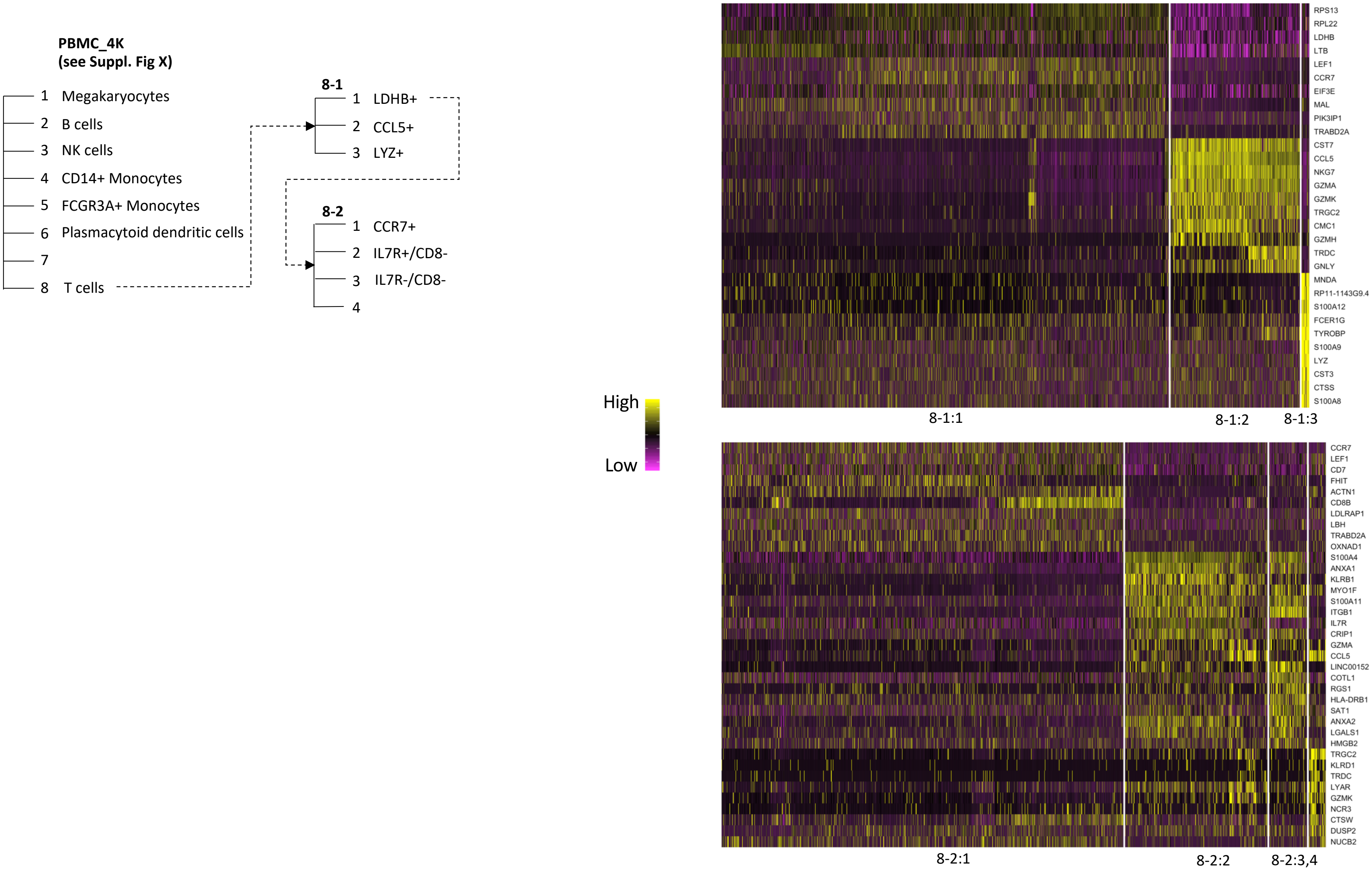

PBMC_4K
(see Suppl. Fig X)
1 Megakaryocytes
2 B cells
3 NK cells
4 CD14+ Monocytes
5 FCGR3A+ Monocytes
6 Plasmacytoid dendritic cells
7
8 T cells
8-1
1 LDHB+
2 CCL5+
3 LYZ+
8-2
1 CCR7+
2 IL7R+/CD8-
3 IL7R-/CD8-
4
High
Low
RPS13
RPL22
LDHB
LTB
LEF1
CCR7
EIF3E
MAL
PIK3IP1
TRABD2A
CST7
CCL5
NKG7
GZMA
GZMK
TRGC2
CMC1
GZMH
TRDC
GNLY
MNDA
RP11-1143G9.4
S100A12
FCER1G
TYROBP
S100A9
LYZ
CST3
CTSS
S100A8
8-1:1
8-1:2
8-1:3
CCR7
LEF1
CD7
FHIT
ACTN1
CD8B
LDLRAP1
LBH
TRABD2A
OXNAD1
S100A4
ANXA1
KLRB1
MYO1F
S100A11
ITGB1
IL7R
CRIP1
GZMA
CCL5
LINC00152
COTL1
RGS1
HLA-DRB1
SAT1
ANXA2
LGALS1
HMGB2
TRGC2
KLRD1
TRDC
LYAR
GZMK
NCR3
CTSW
DUSP2
NUCB2
8-2:1
8-2:2
8-2:3,4

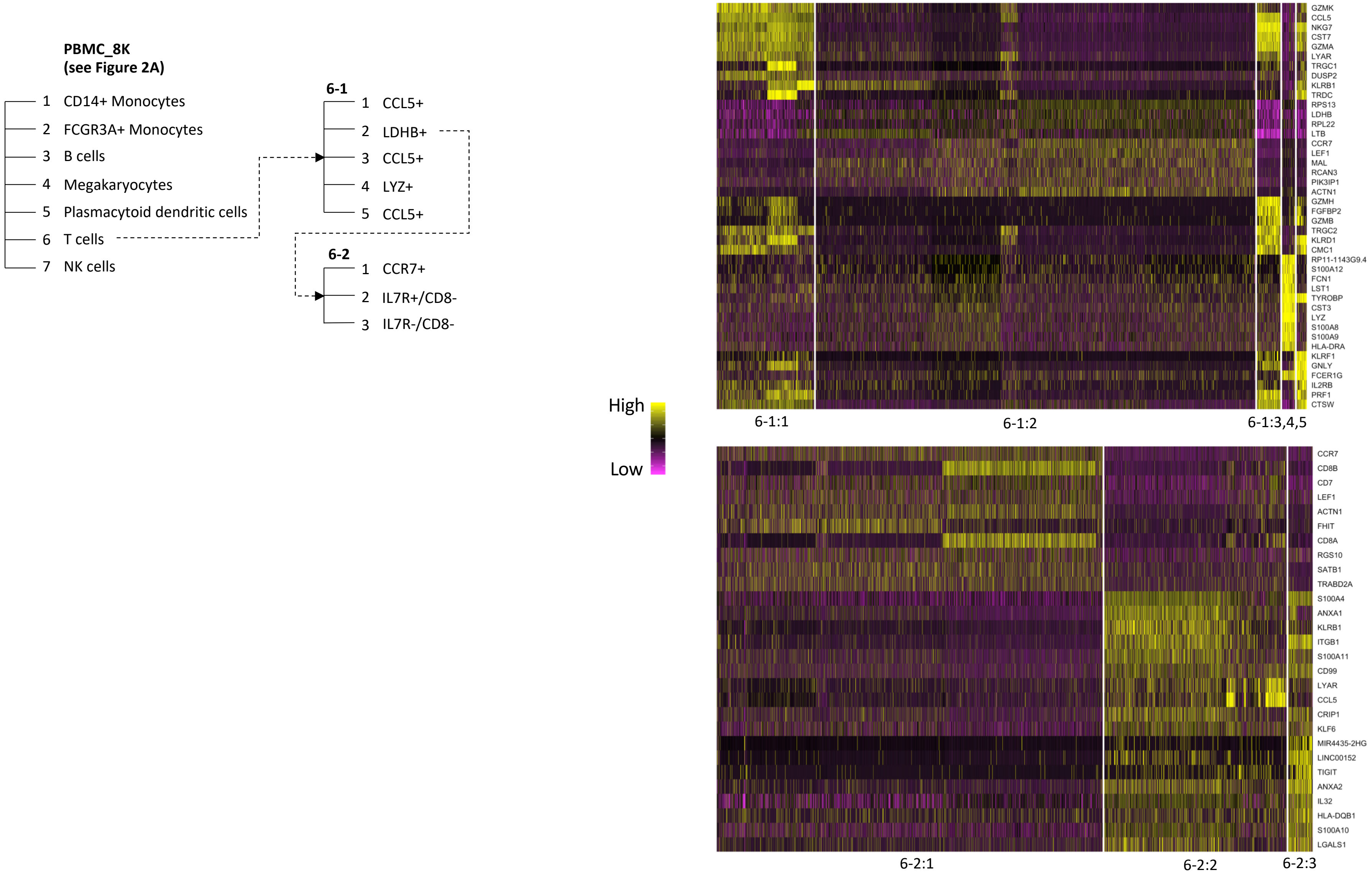
PBMC_8K
(see Figure 2A)
1 CD14+ Monocytes
2 FCGR3A+ Monocytes
3 B cells
4 Megakaryocytes
5 Plasmacytoid dendritic cells
6 T cells
7 NK cells
6-1
1 CCL5+
2 LDHB+
3 CCL5+
4 LYZ+
5 CCL5+
6-2
1 CCR7+
2 IL7R+/CD8-
3 IL7R-/CD8-
High
Low
GZMK
CCL5
NKG7
CST7
GZMA
LYAR
TRGC1
DUSP2
KLRB1
TRDC
RPS13
LDHB
RPL22
LTB
CCR7
LEF1
MAL
RCAN3
PIK3IP1
ACTN1
GZMH
FGFBP2
GZMB
TRGC2
KLRD1
CMC1
RP11-1143G9.4
S100A12
FCN1
LST1
TYROBP
CST3
LYZ
S100A8
S100A9
HLA-DRA
KLRF1
GNLY
FCER1G
IL2RB
PRF1
CTSW
6-1:1
6-1:2
6-1:3,4,5
CCR7
CD8B
CD7
LEF1
ACTN1
FHIT
CD8A
RGS10
SATB1
TRABD2A
S100A4
ANXA1
KLRB1
ITGB1
S100A11
CD99
LYAR
CCL5
CRIP1
KLF6
MIR4435-2HG
LINC00152
TIGIT
ANXA2
IL32
HLA-DQB1
S100A10
LGALS1
6-2:1
6-2:2
6-2:3

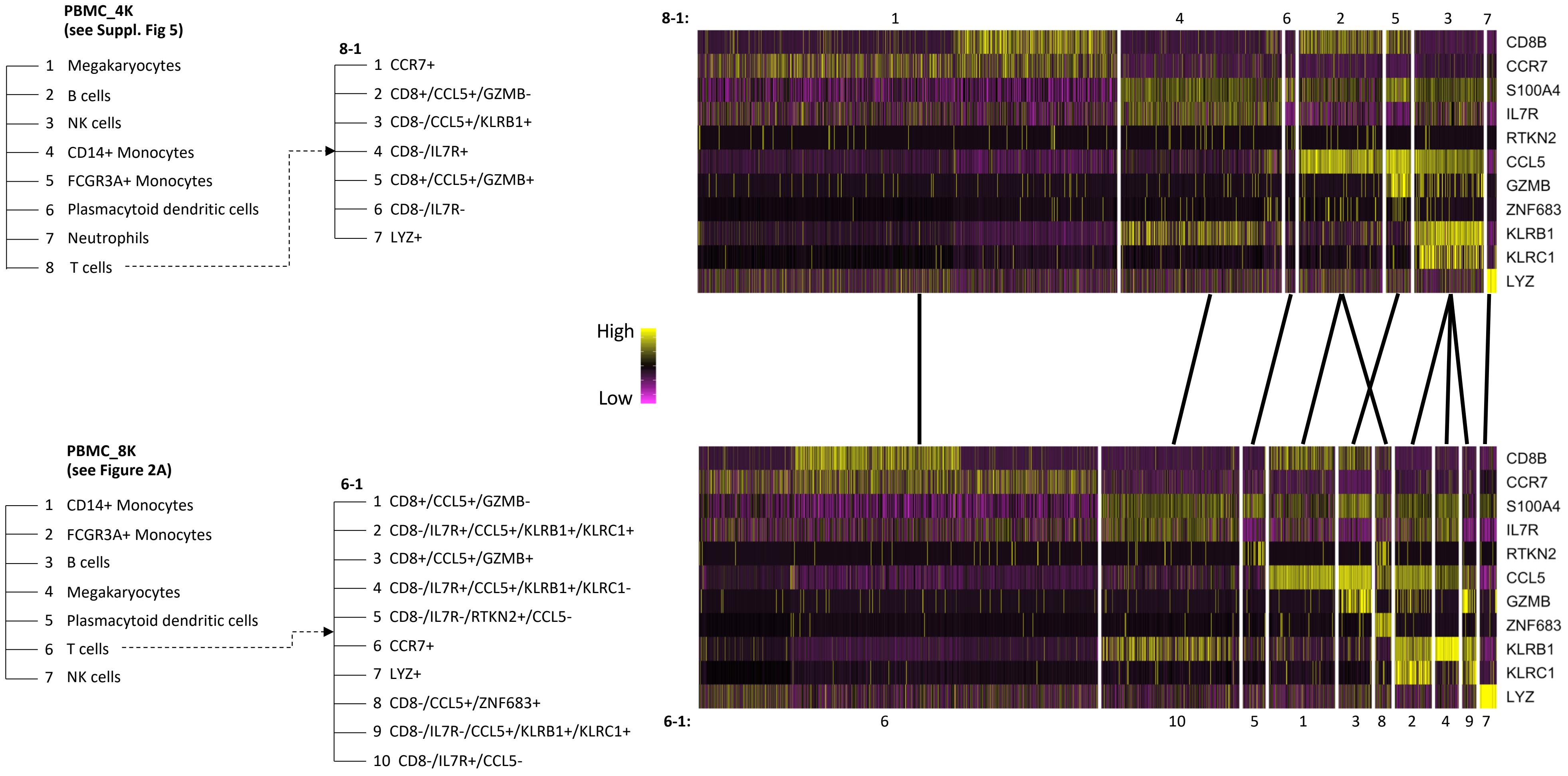

PBMC_4K
(see Suppl. Fig 5)
1 Megakaryocytes
2 B cells
3 NK cells
4 CD14+ Monocytes
5 FCGR3A+ Monocytes
6 Plasmacytoid dendritic cells
7 Neutrophils
8 T cells
8-1
1 CCR7+
2 CD8+/CCL5+/GZMB-
3 CD8-/CCL5+/KLRB1+
4 CD8-/IL7R+
5 CD8+/CCL5+/GZMB+
6 CD8-/IL7R-
7 LYZ+
8-1:
1 4 6 2 5 3 7
CD8B
CCR7
S100A4
IL7R
RTKN2
CCL5
GZMB
ZNF683
KLRB1
KLRC1
LYZ
High
Low
PBMC_8K
(see Figure 2A)
1 CD14+ Monocytes
2 FCGR3A+ Monocytes
3 B cells
4 Megakaryocytes
5 Plasmacytoid dendritic cells
6 T cells
7 NK cells
6-1
1 CD8+/CCL5+/GZMB-
2 CD8-/IL7R+/CCL5+/KLRB1+/KLRC1+
3 CD8+/CCL5+/GZMB+
4 CD8-/IL7R+/CCL5+/KLRB1+/KLRC1-
5 CD8-/IL7R-/RTKN2+/CCL5-
6 CCR7+
7 LYZ+
8 CD8-/CCL5+/ZNF683+
9 CD8-/IL7R-/CCL5+/KLRB1+/KLRC1+
10 CD8-/IL7R+/CCL5-
6-1:
6 10 5 1 3 8 2 4 9 7

PBMC_4K

Expression of top 10 marker genes for 7 T cell subgroups

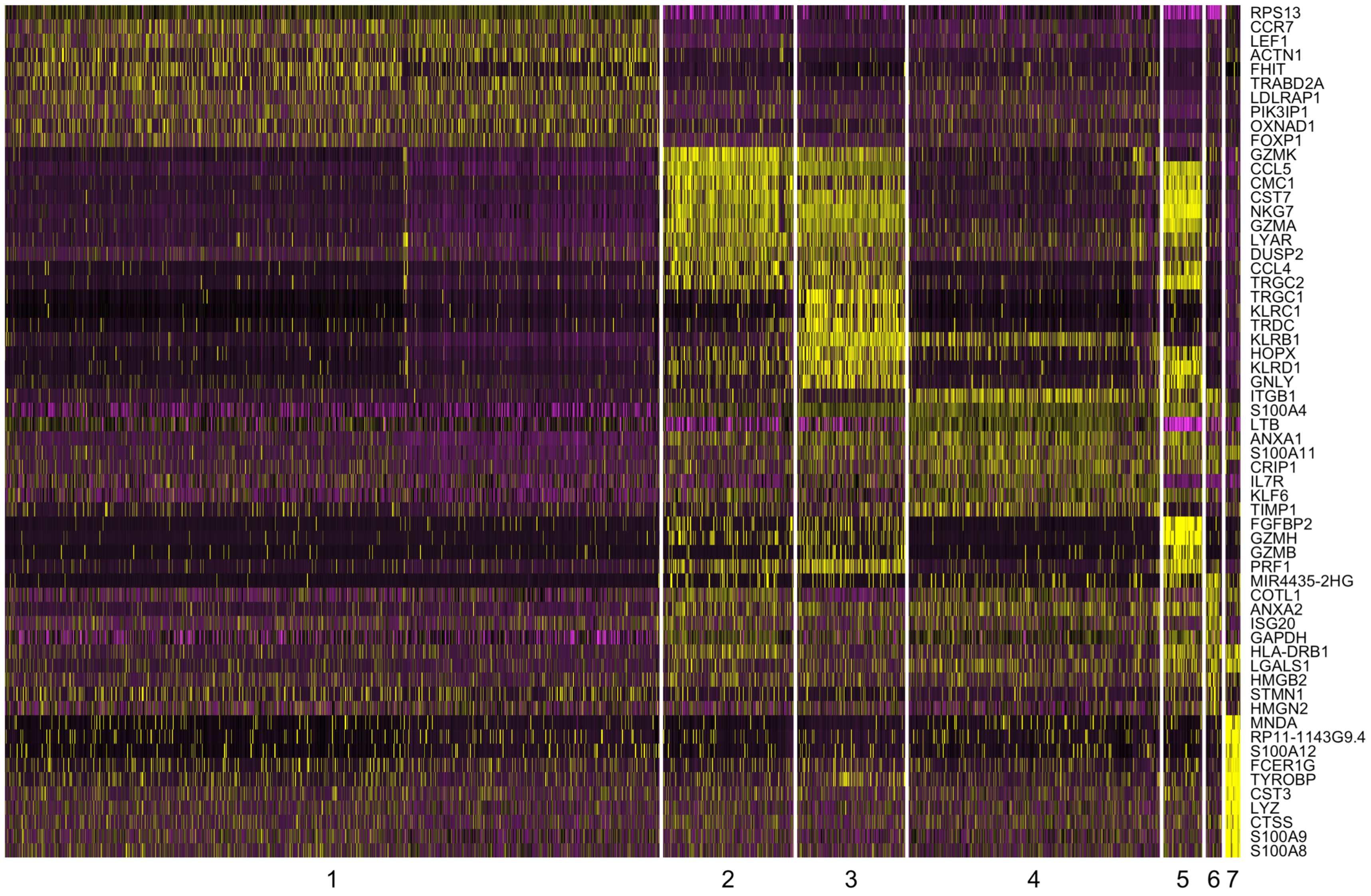

PBMC_8K

Expression of top 10 marker genes for 10 T cell subgroups

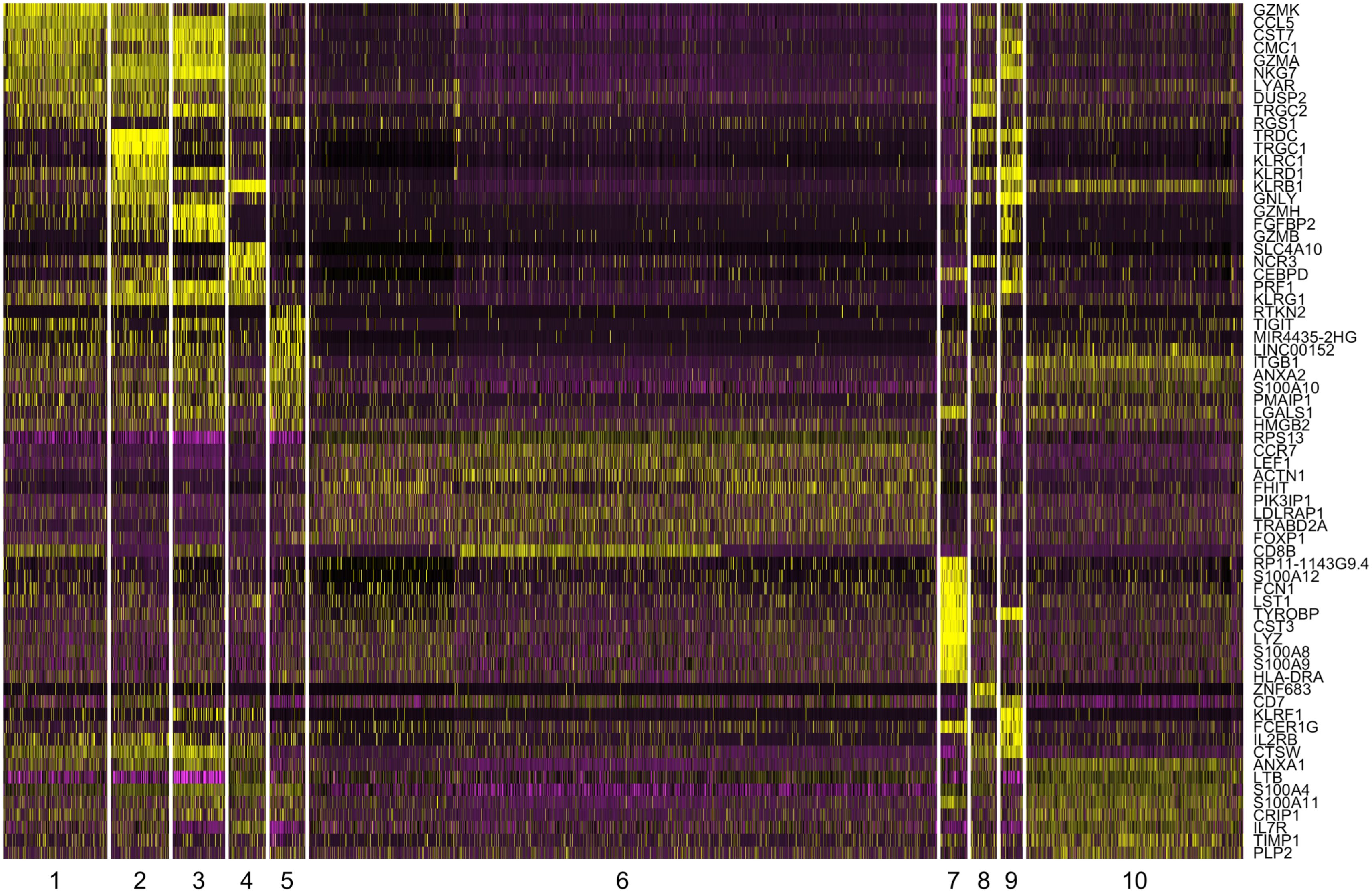

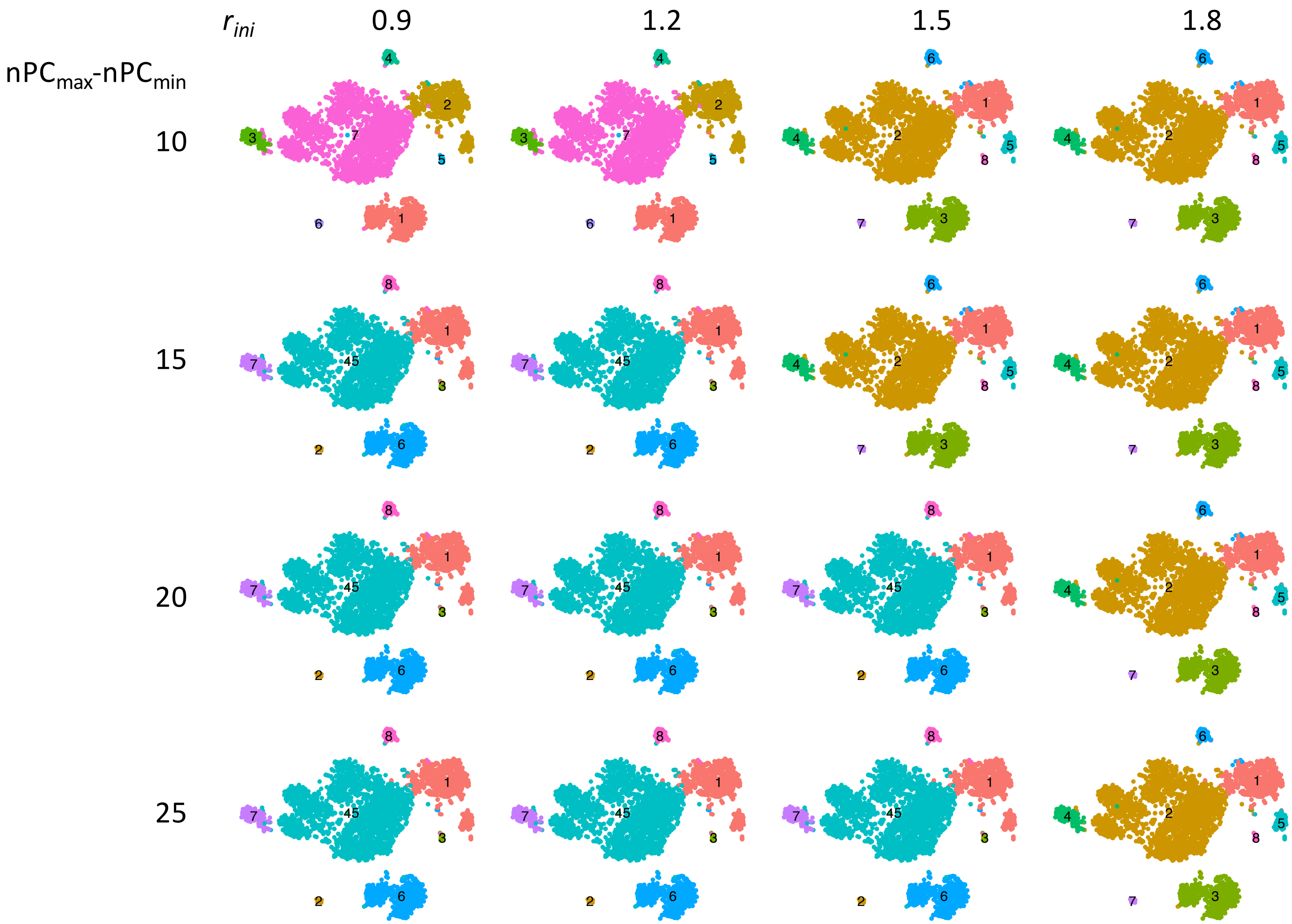

$r_{ini}$
0.9
1.2
1.5
1.8
$nPC_{max}$-$nPC_{min}$
10
15
20
25

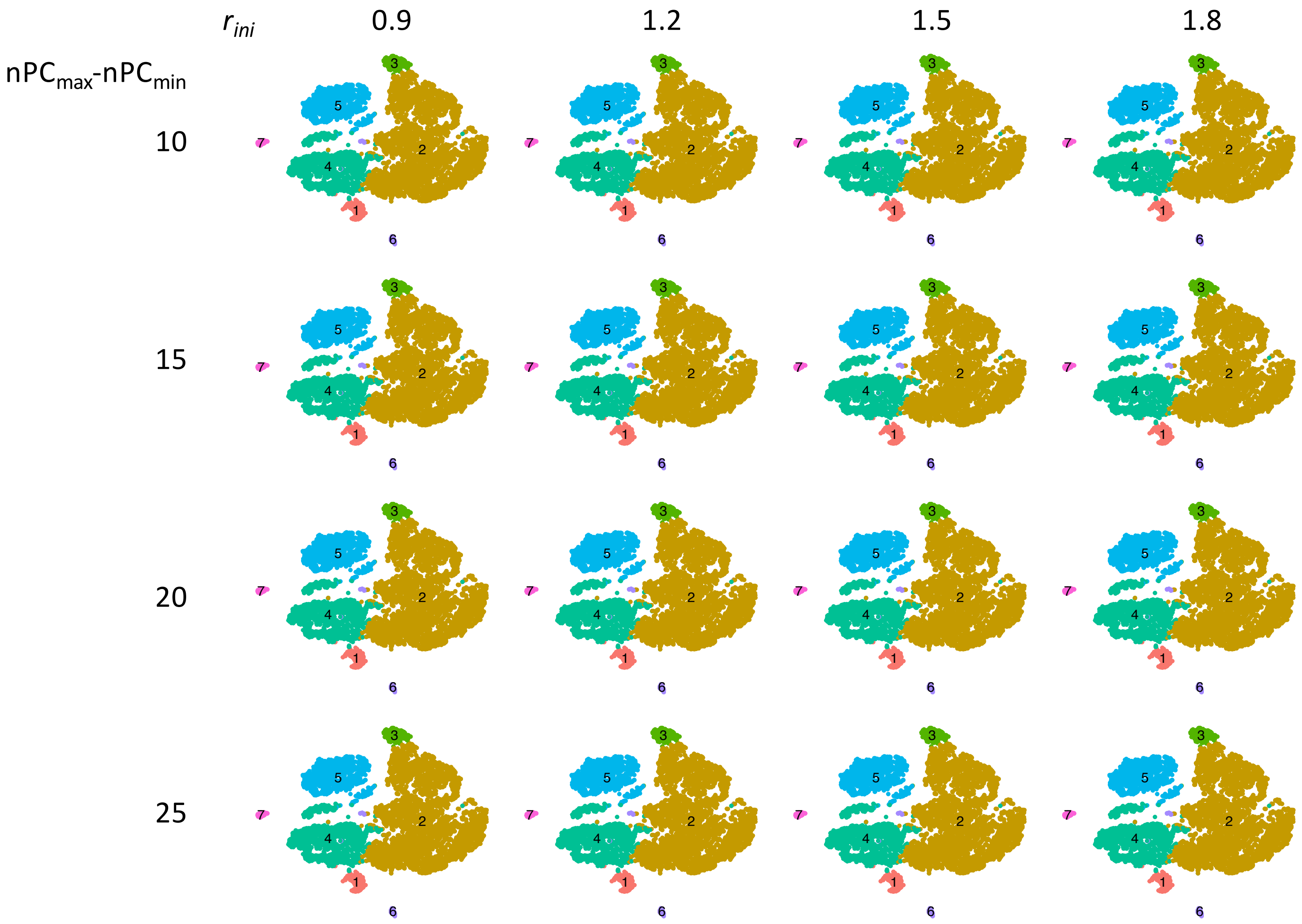

$r_{ini}$
0.9
1.2
1.5
1.8
$nPC_{max}$-$nPC_{min}$
10
15
20
25

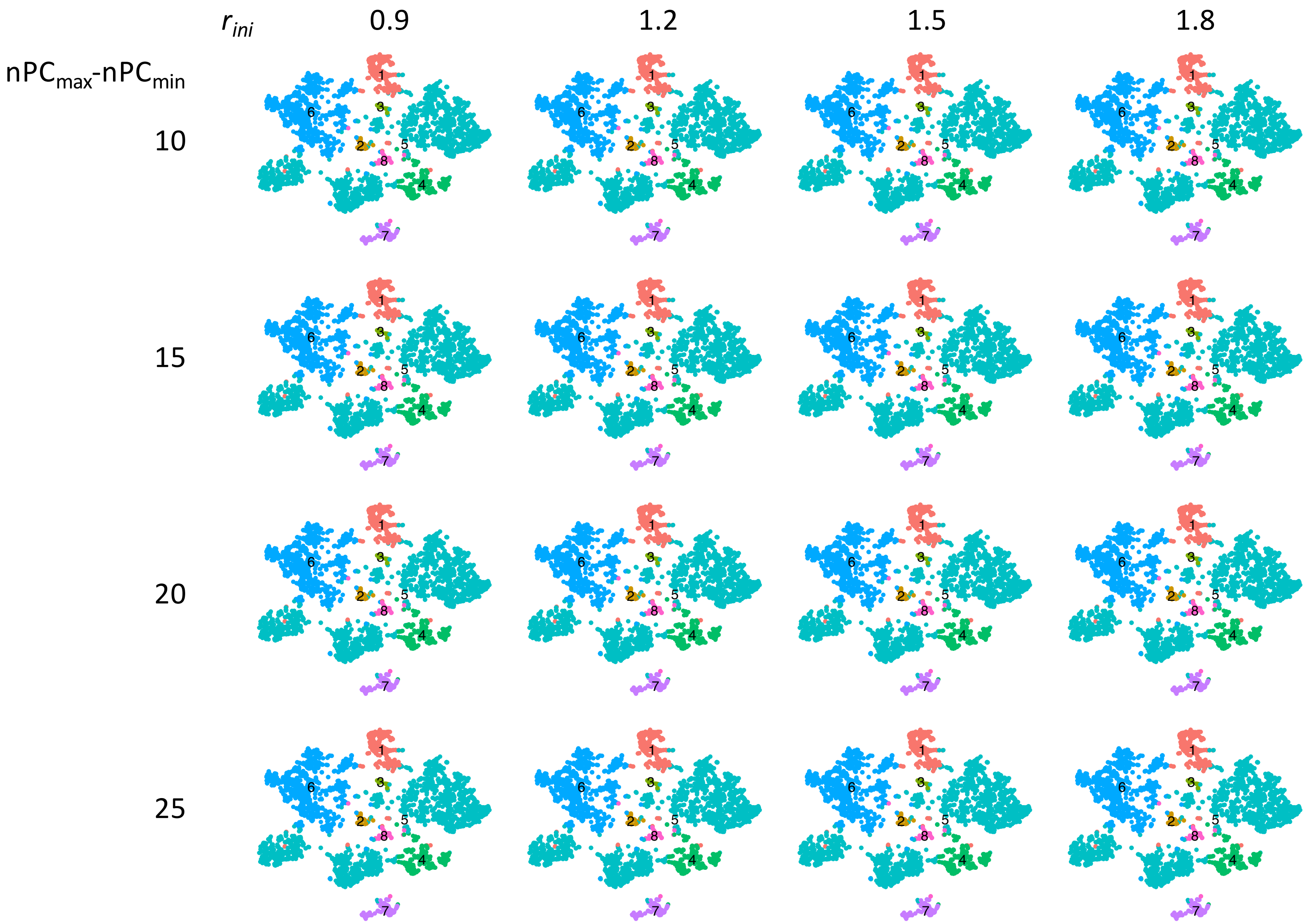

$r_{ini}$
0.9
1.2
1.5
1.8
$nPC_{max}$-$nPC_{min}$
10
15
20
25

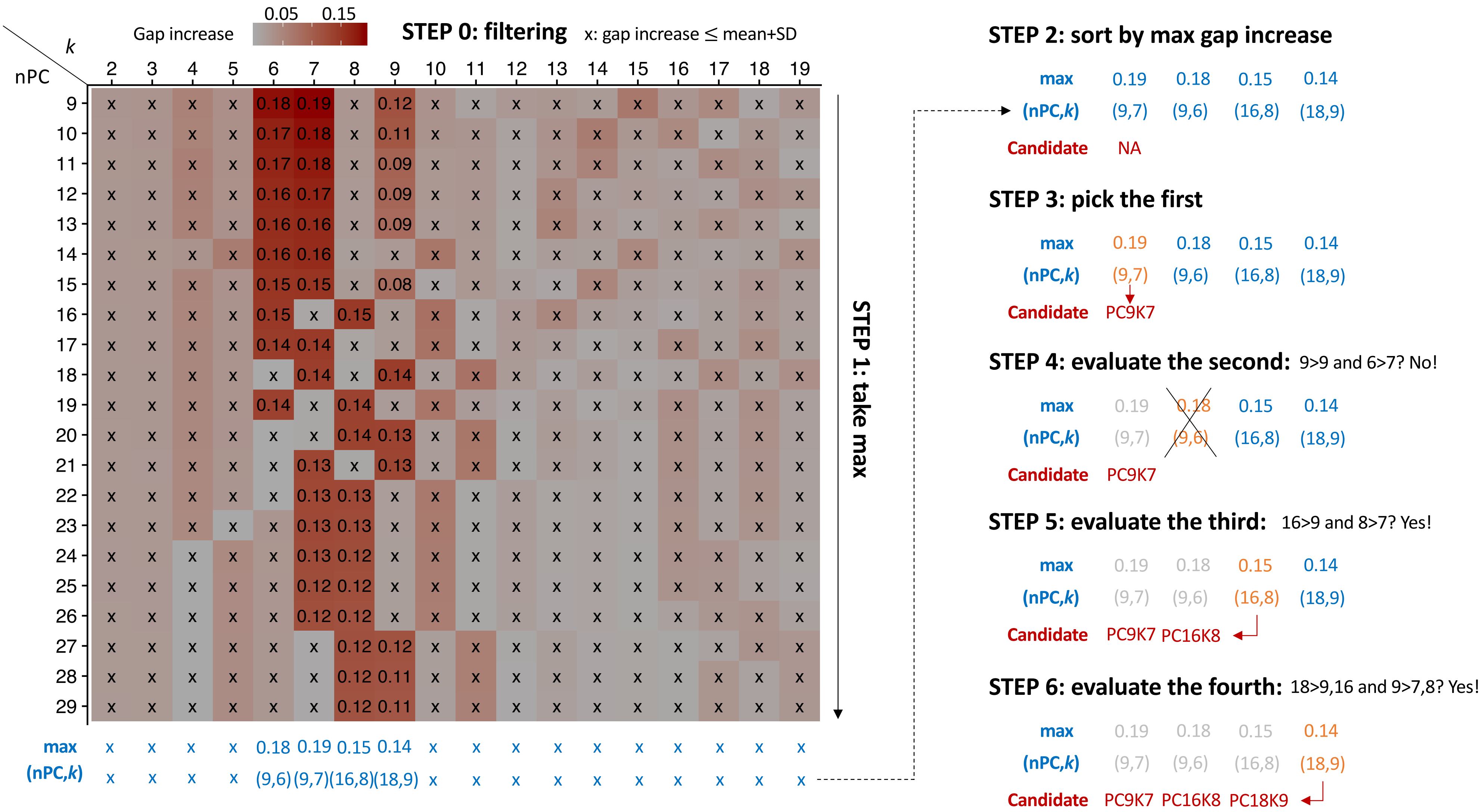
Gap increase
0.05 0.15
STEP 0: filtering
x: gap increase ≤ mean+SD
k
nPC
STEP 1: take max
max x x x x 0.18 0.19 0.15 0.14 x x x x x x x x x x
(nPC,k) x x x x (9,6) (9,7) (16,8) (18,9) x x x x x x x x x x
STEP 2: sort by max gap increase
max 0.19 0.18 0.15 0.14
(nPC,k) (9,7) (9,6) (16,8) (18,9)
Candidate NA
STEP 3: pick the first
max 0.19 0.18 0.15 0.14
(nPC,k) (9,7) (9,6) (16,8) (18,9)
Candidate PC9K7
STEP 4: evaluate the second: 9>9 and 6>7? No!
max 0.19 0.18 0.15 0.14
(nPC,k) (9,7) (9,6) (16,8) (18,9)
Candidate PC9K7
STEP 5: evaluate the third: 16>9 and 8>7? Yes!
max 0.19 0.18 0.15 0.14
(nPC,k) (9,7) (9,6) (16,8) (18,9)
Candidate PC9K7 PC16K8
STEP 6: evaluate the fourth: 18>9,16 and 9>7,8? Yes!
max 0.19 0.18 0.15 0.14
(nPC,k) (9,7) (9,6) (16,8) (18,9)
Candidate PC9K7 PC16K8 PC18K9
